# Clonal Level Lineage Commitment Pathways of Hematopoietic Stem Cells *In Vivo*

**DOI:** 10.1101/262774

**Authors:** Rong Lu, Agnieszka Czechowicz, Jun Seita, Du Jiang, Irving L. Weissman

## Abstract

While hematopoietic stem cells (HSCs) have been extensively studied at the population level, little is known about the lineage commitment of individual clones. Here, we provide comprehensive maps of *in vivo* HSC clonal development in mice under homeostasis and after depletion of the endogenous hematopoietic system. Under homeostasis, all donor-derived HSC clones regenerate blood homogeneously throughout all measured stages and lineages of hematopoiesis. In contrast, after the hematopoietic system has been depleted by irradiation or by an anti-ckit antibody, only a small fraction of donor-derived HSC clones differentiates while dominantly expanding and exhibiting lineage bias. We identified the cellular origins of clonal dominance and lineage bias, and uncovered the lineage commitment pathways that lead HSC clones to differential blood production. This study reveals surprising alterations in HSC regulation by irradiation, and identifies the key hematopoiesis stages that may be manipulated to control blood production and balance.

**SIGNIFICANCE STATEMENT:** Hematopoietic stem cells (HSCs) sustain daily blood production through a complex step-wise lineage commitment process. In this work, we present the first comprehensive study of HSC lineage commitment at the clonal level and identify new HSC regulatory mechanisms that are undetectable by conventional population level studies. First, we uncover distinct HSC clonal pathways that lead to differential blood production and imbalances. Second, we reveal that HSC regulation under physiological conditions is strikingly different from that after injury. Third, we present a comprehensive map of HSC activities in vivo at the clonal level.

## Introduction

Hematopoietic stem cells (HSCs) sustain the blood and immune systems through a complex lineage commitment process (1–3). This process involves several steps during which HSCs become progressively more specified in their potential and eventually give rise to mature blood and immune cells with distinct functions. This step-wise lineage commitment forms the basis of the hematopoietic hierarchy and establishes a paradigm for studying cellular development, differentiation, and malignancy. While the hematopoietic hierarchy has been extensively studied to describe the aggregate differentiation process of the HSC population, little is known about the lineage commitment of individual HSC clones.

Knowledge of HSC clonal level lineage commitment can reveal new insights into HSC regulatory mechanisms. Many regulatory factors act on individual HSC clones and not on the population as a whole. For example, the HSC regulatory microenvironment, HSC niche, may not affect all HSCs in an organism equally, and may instead act directly on resident HSC clones through direct contact or through the tuning of local cytokine concentrations (4–7). The distinct characteristics of HSC clones have also been inferred by several recent studies on the clonality of blood cells, suggesting that HSC clones are heterogeneous and possess differentiation preferences towards either myeloid or lymphoid lineages (8–19). Although such lineage bias is averaged at the population level, it plays important roles in aging, immune deficiency, and many hematopoietic disorders involving an unbalanced hematopoietic system (8, 14, 20–23). The existence of lineage bias indicates that HSC differentiation at the population level is an amalgamation of diverse lineage commitments of individual HSC clones. Disentangling the heterogeneous hematopoiesis is not only essential for understanding HSC regulatory mechanisms, but may also provide new insights into the origin of hematological diseases and may identify new therapeutic targets.

Virtually all HSC studies rely on irradiation-mediated transplantation, including those suggesting HSC lineage bias (8, 10, 12, 13, 15). HSCs are usually purified using cell surface markers *ex vivo* and then transplanted into a different host for study, where the activities of donor HSCs can be distinguished from those of other cells (1–3, 24). The transplantation procedure is almost always accompanied by pre-conditioning the recipient with irradiation (1–3, 24). Irradiation is applied to enhance donor HSC engraftment by massively depleting the recipient’s endogenous hematopoietic and blood cells (25). It is also widely used in the clinical treatment of cancers and hematopoietic disorders to eliminate diseased cells. An alternative pre-transplantation conditioning regimen to irradiation has recently been developed using an anti-ckit antibody, named ACK2 (26). ACK2 selectively eradicates hematopoietic stem and progenitor cells while leaving mature hematopoietic cells intact. This targeted regimen perturbs the hematopoietic system to a lesser degree than irradiation.

While pre-conditioning of the recipient is necessary to obtain high levels of HSC engraftment, all conditioning regimens, to various degrees, injure and derange the HSC niches that normally regulate HSCs (27, 28). Although damaged niches can be restored to some extent after conditioning, it is unclear whether HSC regulation in restored niches still resembles that under normal physiological conditions. For instance, the fraction of HSCs in the cell cycle is significantly higher in irradiated mice than in untreated mice (29). Additionally, recent studies of hair follicle stem cells and intestinal stem cells suggest that tissue repair and homeostasis may be sustained by distinct stem cells and through different mechanisms (30, 31). Studying HSC lineage commitment absent irradiation conditioning is crucial for understanding natural HSC function.

HSCs can be transplanted without the use of conditioning (26, 32) by taking advantage of natural HSC migration in the peripheral blood (33–35). After unconditioned transplantation, donor HSCs injected into the peripheral blood numerically outcompete endogenous migrating HSCs in homing into available niches and subsequently participate in normal hematopoiesis for the rest of the organism’s lifetime. Unconditioned transplantation minimally perturbs the natural hematopoiesis and provides an optimal experimental model for studying homeostatic hematopoiesis of HSCs.

Unfortunately, unconditioned transplantation produces low engraftment rates even after repeat transplantations (26, 32). Tracking the low number of individual HSCs *in vivo* that engraft after unconditioned transplantation had been technically prohibitive until the recent development of an *in vivo* clonal tracking technology (36–38). By combining genetic barcoding and high-throughput sequencing, this technology offers both the high throughput and the high sensitivity necessary for tracking the low number of HSCs that engraft in unconditioned transplantation. It allows for the direct examination of the clonality of HSCs and other hematopoietic progenitors, providing a clonal view of the entire hematopoietic process for the first time. This technology also offers the capacity to simultaneously track many HSCs in a single mouse, and thereby reveals the interactions of different HSC clones in the same host.

Using this clonal tracking technology (36), we examined clonal level HSC development throughout multiple stages of lineage commitment in the absence of conditioning and after conditioning with lethal irradiation or ACK2 treatment. These data provides a comprehensive view of HSC activities at the clonal level and reveals the underlying lineage commitment pathways of HSC clones. The discovery of these clonal pathways provides a new interpretation of studies using irradiation-mediated transplantation and identifies key stages of HSC development that may serve as potential targets for the treatment of different hematopoietic disorders.

## RESULTS

Purified HSCs were genetically labeled with unique 33-bp barcodes at the single cell level using a lentiviral vector prior to transplantation (Fig. 1A). The genetic barcode is inherited by every progeny of a barcoded HSC through cell division and development. Thus, the viral labeling establishes a one-to-one mapping of a single HSC clone with a unique barcode (36). Donor-derived HSCs and their progenies were harvested 22 weeks after transplantation, when blood reconstitution had returned to a steady state (2, 3, 10, 24, 39, 40). Barcodes recovered from these hematopoietic populations were identified and quantified using high throughput sequencing.

**Figure 1.**
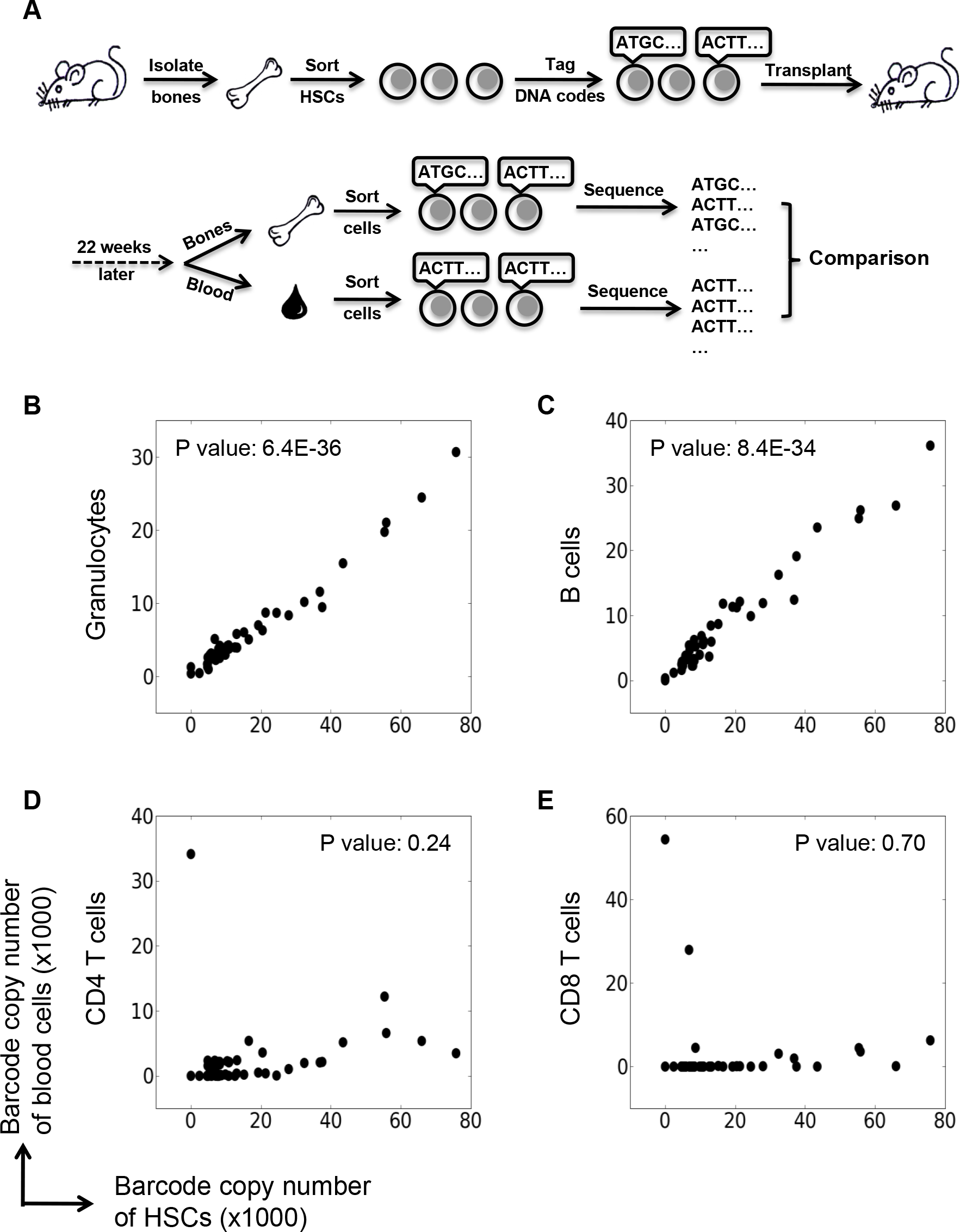
Comparing Clonality of Hematopoietic Stem Cells (HSCs) with Clonality of Blood Cells. (A) Experimental Design. Donor HSCs are harvested from the bone marrow and genetically labeled with 33bp barcodes using a lentiviral vector. Barcoded HSCs are transplanted into mice pre-conditioned with either irradiation or an anti-ckit antibody, or into mice without any conditioning (unconditioned). 22 weeks after transplantation, donor derived hematopoietic stem/progenitor cells and mature blood cells are isolated from bone marrow and peripheral blood respectively. Barcodes are extracted and analyzed as described elsewhere (36). (B-E) Barcode copy numbers from HSCs are compared with those from blood cells after unconditioned transplantation. Each dot represents a unique barcode that is used to track a single HSC clone. The x and y axes represent barcode copy numbers of different cell populations. The two-tailed P values of the Pearson correlation are shown to quantify the significance of the linear correlation. These scatter plots depict data from a single representative mouse. Data from all eight mice are shown in Figure S1.

The number of times that a sequencer detects a unique barcode sequence is referred to as the barcode copy number, indicating the abundance of the barcode in a cell population. This number is roughly proportional to the number of cells that carry the barcode. As a donor-derived HSC develops through the various stages of lineage commitment after transplantation, we can obtain the copy numbers of its barcode from different cell populations at the various developmental stages (1–3). For example, the copy numbers of the same barcode can be measured in HSCs, in granulocytes, in B cells and in other cell populations. While the absolute copy number of a barcode from each cell population is influenced by the amount of barcode DNA loaded onto the sequencer and the PCR amplification used to recover the barcode, the ratio of barcode copy numbers across different cell populations is informative. Equal ratios across different cell populations indicate that the respective clones expand at the same rate. On the other hand, dissimilar ratios indicate that some clones expand at a faster rate than the others.

The relative copy numbers of all barcodes from a population represent its “clonal composition”. Comparing the clonal composition across various stages of differentiation, individual HSC clones can be mapped to precise lineages and stages of the hematopoietic hierarchy in a semi-quantitative manner. This reveals the relative contributions of individual HSC clones to different hematopoietic lineages and their relative expansion during the step-wise lineage commitment process. As all cell populations were collected 22 weeks after transplantation, the data represents a snapshot of a dynamic hematopoiesis.

### HSC Clones Homogenously Differentiate After Unconditioned Transplantation

22 weeks after unconditioned transplantation, all barcode copy number ratios of HSCs to granulocytes are approximately equal (Fig. 1B and S1), indicating that every HSC clone expands at a similar rate between these two stages. The same relationship holds true for barcode copy numbers between the HSC and B cell stages (Fig. 1C and S1). Taken together, the data indicates that donor-derived HSC clones homogenously contribute to granulocytes and B cells after unconditioned transplantation. However, not every HSC barcode can be found amongst CD4 T cells or CD8 T cells in the peripheral blood (Fig. 1D–E, and S1), suggesting that some engrafted HSC clones do not contribute to the mature T cell repertoire. This explains why a chronic myelocytic leukemia clone from the HSC pool can be found in granulocytes, monocytes, erythrocytes, and B cells, but is rarely present in T cells (41). It is likely that the migration of the T cell precursors, mainly the common lymphocyte progenitors (CLP), to the thymus or the maturation of T cells within the thymus is episodic and restricts the number of HSC clones that eventually contribute to mature T cells (42). Thus, it is important to use a well-defined blood cell type for HSC clonal tracking studies. In this study we use granulocytes and B cells, both of which mature in the bone marrow, to analyze myeloid versus lymphoid lineage bias.

To determine how the clonal contribution of different HSCs evolves through the multiple stages of hematopoiesis, we examined the intermediate progenitors of myeloid and lymphoid differentiation (Fig. 2A–B and S2) (1–3). After unconditioned transplantation, the relative copy numbers of barcodes in HSCs remain generally constant in the multipotent progenitors, the oligopotent progenitors and the terminally differentiated granulocytes and B cells (Fig. 2A–B and S2). This indicates a homogeneous clonal contribution of donor HSCs to the various stages of hematopoiesis. A few clones expand at the progenitor stages, but this infrequent expansion is not reflected in blood cells (Fig. 2A–B and S2). Therefore, in unconditioned mice, HSC lineage commitment progresses with an equal contribution from each clone.

**Figure 2.**
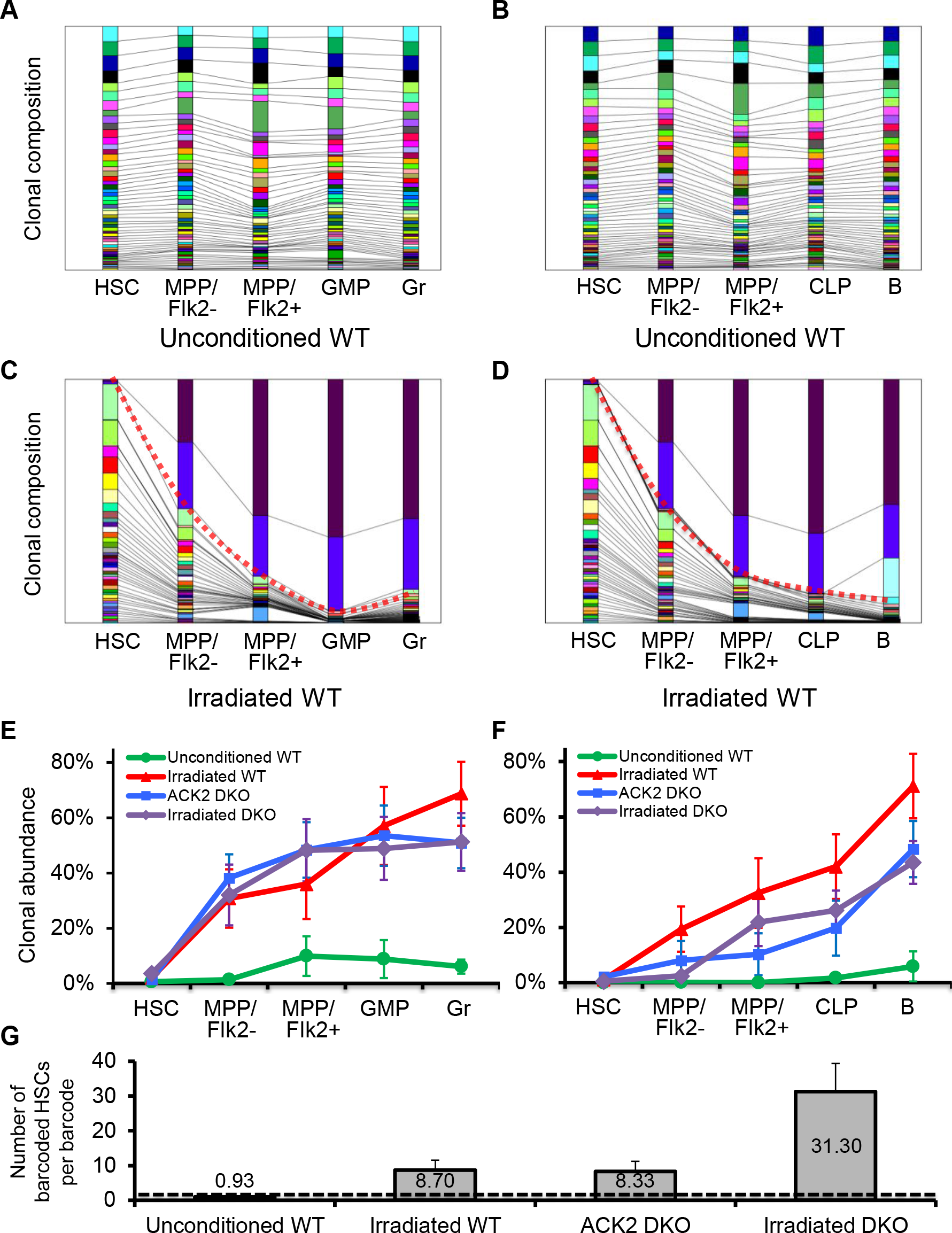
Pre-transplantation Conditioning Induces Dominant Differentiation of HSC Clones. (A-D) Clonal compositions at each stage of HSC differentiation. Each column represents a hematopoietic population. Each colored section in a column represents one distinct genetic barcode, corresponding to a HSC clone. The size of each colored section indicates its relative abundance within each cell population. Identical barcodes from different cell populations (columns) are shown in the same color and are connected by lines. Red dotted lines highlight clones that exhibit dominant differentiation in irradiated mice. Hematopoietic stem cell (HSC), Flk2− multipotent progenitor (MPP^Flk2−^), Flk2+ multipotent progenitor (MPP^Flk2+^), granulocyte/monocytic progenitor (GMP), common lymphocyte progenitor (CLP), granulocyte (Gr) and B cell (B) are arranged along myeloid differentiation stages (A, C) and lymphoid differentiation stages (B, D). Barcodes are arranged from top to bottom according to their abundances in terminally differentiated cells in the right-most column of each panel. Shown are data from a recipient mouse not treated with any pre-transplantation conditioning (A, B) and a recipient mouse treated with lethal irradiation prior to transplantation (C, D). Data from all eight unconditioned mice and seven irradiated mice are shown in Figure S2. (E, F) The percentage of barcodes representing dominant clones at each stage of HSC differentiation under various transplantation conditions. Dominant clones are defined as those whose relative copy numbers in blood cells (granulocytes or B cells) are more than five times their relative copy numbers in HSCs. Similar results are obtained when dominant clones are defined by different threshold values (Figure S3 A-B). WT: wild type mice. DKO: Rag2^−/−^ϒc^−/−^ mice. ACK2: a clone of anti-ckit antibody. (G) Number of harvested barcoded HSCs carrying the same barcode 22 weeks after transplantation. GFP+ HSCs are counted as barcoded HSCs at the time of harvest, as GFP is constantly expressed in the barcode vectors. The dashed line shows the initial HSC to barcode ratio of 1 at the time of transplantation. (E-G) Error bars show the standard errors of the means for all mice under the same transplantation conditions.

### Pre-transplantation Conditioning Induces Dominant Differentiation of HSC Clones

To compare HSC lineage commitment after unconditioned transplantation versus that after conditioned transplantation, identical numbers of barcoded HSCs were transplanted into unconditioned, lethally irradiated, or ACK2-treated mice. As ACK2 effectively facilitates HSC engraftment only in Rag2^−/−^ϒc^−/−^ double knockout (DKO) mice, control transplantations using irradiation were also performed on the DKO mice. In contrast to the homogeneous differentiation of all engrafted HSC clones in unconditioned mice (Fig. 2A–B and S2), in irradiated mice a small fraction of engrafted HSC clones expands substantially faster than other clones during differentiation and supplies the majority of granulocytes and B cells (Fig. 2C–D and S2). We call this clonal behavior “dominant differentiation” and the clones that exhibit this behavior “dominant”. It is important to note that dominant clones in irradiated mice are not dominant at the HSC stage, but only become dominant at the intermediate progenitor stages as measured at week 22 post transplantation (Fig. 2C–D and S2). In a conditioned mouse, more than half of the measured granulocytes and B cells descend from the dominant differentiation of a few HSC clones (Fig. 2E–F and S3A–B). The dominant differentiation of HSC clones is present at similar levels after irradiation and ACK2 treatment (Fig. 2E–F and S3A–B), indicating that it is not specific to either regimen.

While pre-transplantation conditionings induce dominant clonal expansion in HSC differentiation, we ask whether it also influences HSC self-renewal. At the time of transplantation, each barcode labels one HSC (36). If self-renewed, this HSC becomes multiple HSCs that all carry identical barcodes. Thus, the ratio of the number of HSCs to the number of unique barcodes increases with self-renewal. We found that after unconditioned transplantation, each barcode is derived from about one barcoded HSC (Fig. 2G). This is very similar to the cell-to-barcode ratio of the original HSC infection (36). Therefore, donor-derived HSCs have not been significantly amplified nor reduced in recipient mice after unconditioned transplantation. This indicates that HSCs are not pressured to expand during homeostatic hematopoiesis after unconditioned transplantation. In contrast, when the endogenous HSCs have been depleted by transplantation conditioning, donor HSCs must expand via self-renewal to reconstitute the entire HSC pool. Consistent with this prediction, after conditioned transplantation, each barcode is derived from on average more than eight barcoded HSCs (Fig. 2G). Thus, HSCs have experienced at least three cell cycles of self-renewal by week 22 after conditioned transplantation. Therefore, HSCs dominantly expand during self-renewal and differentiation in conditioned mice but not in unconditioned mice (Fig. 2).

### Pre-transplantation Conditioning Induces HSC Lineage Bias

The lineage bias of an HSC clone is determined by its relative contribution to myeloid versus lymphoid lineages. For example, HSC barcodes with myeloid bias would have relatively high copy numbers in myeloid cell types such as granulocytes and relatively low copy numbers in lymphoid cell types such as B cells. After irradiation-mediated transplantation, donor-derived HSC clones can be separated into three groups using the ratio of granulocyte barcode copy numbers to B cell barcode copy numbers (Fig. 3B, 3D, and S4). These three groups represent myeloid bias, lymphoid bias and lineage balance, consistent with previous studies (8, 10, 12, 13, 15). In ACK2-treated mice, HSC clones also exhibit both lineage biases and lineage balance, forming three clearly separated groups (Fig. 3C and S4). The fraction of HSCs exhibiting lineage bias and balance is similar after irradiation and after ACK2 treatment (Fig. S3C). However, no lineage bias is observed when pre-transplantation conditioning is absent (Fig. 3A and S4).

**Figure 3.**
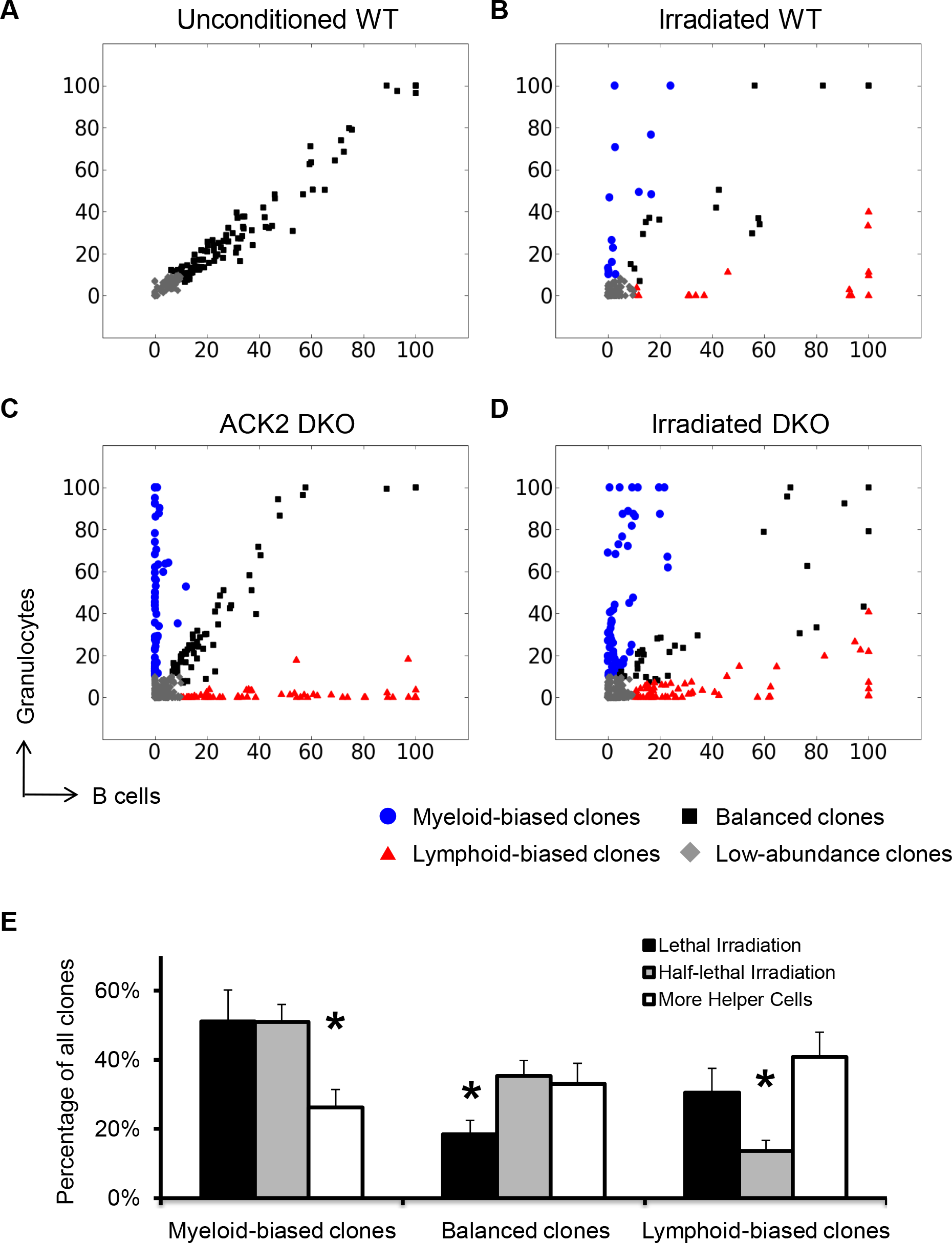
Pre-transplantation Conditioning Induces HSC Lineage Bias. (A-D) Scatter plots comparing barcode copy numbers from granulocytes with barcode copy numbers from B cells in the peripheral blood. Each dot represents a unique barcode that is used to track a single HSC clone. Colors are assigned according to the ratios of the granulocytes barcode copy numbers (myeloid lineage) to B cells barcode copy numbers (lymphoid lineage). Lineage-biased clones are defined as those whose relative copy numbers in one lineage are more than 2.4142 (cotangent 22.5 degree) times their relative copy numbers in the other lineage. Low-abundance clones are excluded from the analysis of lineage bias versus balance. These clones are defined as those whose copy numbers are less than 10% of the maximum copy numbers in both lineages. Shown are the barcodes from all mice examined under each condition. Barcode copy numbers are normalized to the most abundant clone in each cell population of each mouse. Mice with insufficient reads (maximum barcode copy number < 5000) or with a low number of barcodes (less than 3 unique barcodes present in granulocytes and B cells) are excluded from these plots. Raw data from all mice are shown individually in Figure S4. The percentages of barcodes with distinct lineage bias and balance are summarized in Figure S3C. (E) Lineage bias and balance of donor HSCs after various conditions of irradiation-mediated transplantations. Shown are percentages of clones with distinct lineage bias or balance. Lethal irradiation uses 950 cGy and 0.5 million helper cells (whole bone marrow cells). Half lethal irradiation uses 475 cGy and 0.5 million helper cells. The “more helper cells” condition uses 5 million helper cells and 950 cGy. Error bars show the standard errors of the means for all mice under the same transplantation conditions. * P<0.05 by Student’s t-test.

The lineage bias and balance of engrafted clones are also affected by the irradiation dosage and by the number of helper cells used in the transplantation procedure (Fig. 3E). Compared to lethal irradiation, increasing the number of helper cells results in significantly less myeloid-biased clones (Fig. 3E). On the other hand, half-lethal irradiation produces significantly less lymphoid-biased clones (Fig. 3E). The use of more helper cells and the reduction of the irradiation dosage both generate significantly more balanced HSC clones (Fig. 3E). Thus, the observed lineage bias of donor HSCs is highly sensitive to transplantation conditions and is absent when no conditioning regimen is applied (Fig. 3).

### Dominant Differentiation and Lineage Bias Are Connected

As dominant differentiation and lineage bias are both present in conditioned recipients and both absent in unconditioned recipients, we wonder whether they are associated with each other and simultaneously affect the same HSC clones. We separate all HSC clones in conditioned recipients into dominant and non-dominant groups based on their expansion between HSCs and blood cells (including both granulocytes and B cells). We then examine the lineage bias and balance of each group. In irradiated mice, both dominant clones and non-dominant clones exhibit similar proportions of lineage bias and balance (Fig. 4A, 4C and S5). However, in ACK2-treated mice, dominant clones exhibit lineage bias, whereas non-dominant clones exhibit lineage balance (Fig. 4B and S5). In these mice, if dominant clones are excluded, the remaining clones are mostly balanced, as if they had been transplanted without any conditioning (Fig. 4D). This suggests that HSC differentiation regulatory mechanisms active under normal homeostatic conditions are still active in ACK2-treated mice and regulate a subset of engrafted HSC clones. However, these mechanisms are inactivated in irradiated mice. The concurrence of dominant differentiation and lineage bias in HSC clones of ACK2-treated mice indicates a connection between these two phenotypes (Fig. 4B and S5).

**Figure 4.**
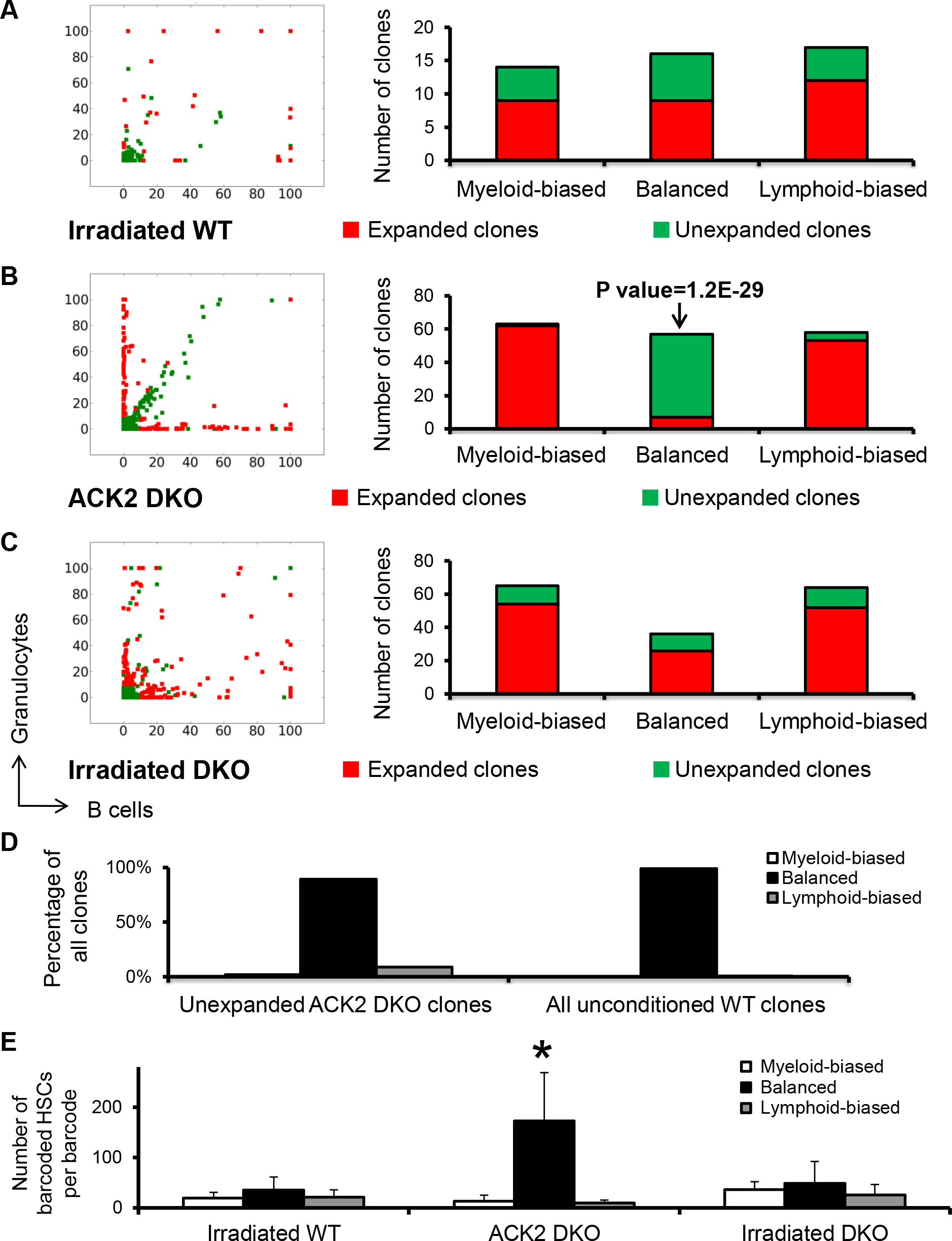
Connections Between Dominant Differentiation and Lineage Bias. (A-C) Dominant clones and non-dominant clones are compared by their lineage bias and balance. (A-C, Left) Plots are generated as described in Figure 3 A-D’s legend. Colors are assigned according to the ratios of HSC barcode copy numbers to granulocytes and B cells copy numbers. Dominant clones are defined as those whose relative copy numbers in blood cells (granulocytes or B cells) are more than five times their relative copy numbers in HSCs. Similar results are obtained when dominant clones are defined by different threshold values (Figure S5). (A-C, Right) The number of clones with lineage bias or balance from all mice under the indicated transplantation condition. P value depicts the probability that a given result is caused by dominant or non-dominant clones randomly becoming lineage-biased or balanced. (D) Lineage bias and balance of non-dominant clones in ACK2-treated Rag2^−/−^ϒc^−/−^ mice (DKO) compared with all clones in unconditioned wild type mice. (E) Numbers of lineage-biased or balanced barcoded HSCs carrying the same barcode. Data are normalized by lymphoid-biased barcodes of each transplantation condition. GFP+ HSCs are counted as barcoded HSCs at the time of harvest, as GFP is constantly expressed in the barcode vectors. Error bars show the standard errors of the means for all mice under the same transplantation conditions. * P<0.05 by Student’s t-test, *** P<0.001.

Lineage bias is associated with clonal expansion not only during HSC differentiation (Fig. 4B) but also during HSC self-renewal (Fig. 4E). In ACK2-treated mice, balanced HSC clones have undergone significantly more self-renewal than other clones in the same mice and more than the clones in irradiated mice during the first 22 weeks after transplantation (Fig. 4E). This suggests that balanced clones preferentially self-renew in ACK2-treated mice. Thus a clonal competition appears to exist in these mice where lineage-balanced clones outcompete lineage-biased clones in numbers at the HSC stage. Taken together, these data suggest a connection between lineage bias, lineage commitment and self-renewal in ACK2-treated mice, where lineage-biased clones dominate in lineage commitment and exhibit low self-renewal (Fig. 4).

### Lineage Bias Arises From Dominant Differentiation at Distinct Lineage Commitment Steps

As lineage bias is associated with dominant differentiation (Fig. 4 and S5), we ask whether lineage bias is derived from the dominant differentiation of any particular lineage commitment steps. We examined the percentage of barcodes representing myeloid-biased clones at various stages of myeloid differentiation and found that the greatest increase in myeloid-biased barcodes occurred between the HSC and Flk2− multipotent progenitor (MPP^Flk2−^) stages (Fig. 5A and S6). In contrast, lymphoid-biased clones undergo the greatest expansion at the last lineage commitment step from CLP to B cell (Fig. 5A and S6). These patterns are found in both ACK2-treated mice and irradiated mice (Fig. 5A). In addition, if we examine all clones that expand dominantly at the CLP-to-B-cell step, they are significantly more likely to end up with lymphoid bias in both ACK2-treated mice and irradiated mice (Fig. 5B). On the other hand, clones that expand dominantly at the HSC-to-MPP^Flk2−^ step are significantly more likely to develop myeloid bias in ACK2-treated mice (Fig. 5B). However, in irradiated mice, these clones could become either myeloid-biased or balanced (Fig. 5B). This indicates that balanced clones in irradiated mice also dominantly differentiate, which will be discussed in depth later (Fig. 6A–B). Taken together, these data suggest that myeloid versus lymphoid lineage bias arises from dominant differentiation at distinct lineage commitment steps (Fig. 5A–B).

**Figure 5.**
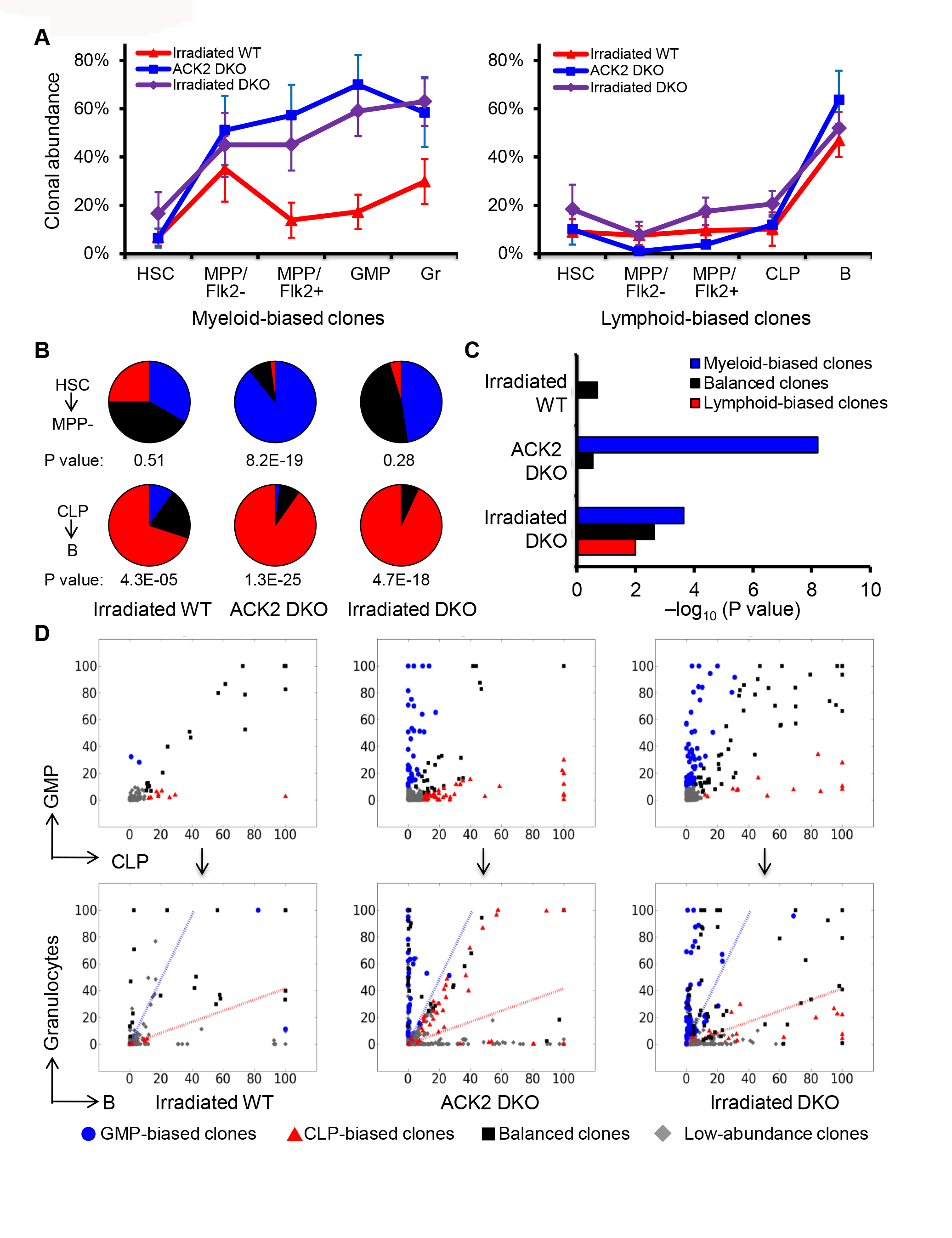
Lineage Bias of HSCs Is Derived From Dominant Differentiation at Distinct Lineage Commitment Steps. (A) Plots were generated as described in Figure 2 E-F’s legend, except that only myeloid-biased clones (left) or lymphoid-biased clones (right) are plotted. Other clones are shown in Figure S6. (B) Pie charts illustrate the lineage bias and balance of the clones that expand dominantly from HSC to MPP^Flk2−^ (upper panels) or from CLP to B cells (lower panels). P value depicts the significance that the clones that dominantly expand during HSC-to-MPP^Flk2−^ commitment become myeloid-biased (upper panel) and that the clones that dominantly expand during CLP-to-B-cell commitment become lymphoid-biased (lower panel). (C) P value depicts the significance that the lineage bias and balance at the progenitor stages is reflected in blood cells. (B-C) P value is calculated based on the assumption that the clones are randomly distributed among the different categories of lineage bias and balance. (D) Plots are generated as described in Figure 3 A-D’s legend. Dot colors in both upper and lower panels are assigned based on the bias between GMPs and CLPs. The dotted lines in the lower panels show the boundary of lineage bias versus balance in blood cells. Analysis based on the lineage bias exhibited in blood cells is shown in Figure S7, where dot colors are assigned based on the bias between granulocytes and B cells.

If myeloid bias arises at the first differentiation step and lymphoid bias arises at the last differentiation step as in the case of ACK2-treated mice (Fig. 5A–B), then myeloid bias but not lymphoid bias should characterize the intermediate progenitor stages. This is validated by data from granulocyte/monocytic progenitors (GMPs) and common lymphocyte progenitors (CLPs) (Fig. 5C–D). In ACK2-treated mice, clones with myeloid bias at the progenitor stages significantly preserve their myeloid bias in blood cells (Fig. 5C–D). Moreover, lymphoid-biased clones and balanced clones at the progenitor stages do not preserve their bias and balance in blood cells, which is consistent with the prediction (Fig. 5C–D). In irradiated wild type mice, all of the clones appear to be balanced at the progenitor stages (Fig. 5D). In contrast, when Rag2^−/−^ϒc^−/−^ (DKO) mice lacking mature lymphoid cells are used as irradiation recipients, the progenitors exhibit lineage bias (Fig. 5D). This bias is moderately preserved in blood cells (Fig. 5C), which will be discussed in depth later (Fig. 6E–F). These data suggest that when transplantation recipients are lineage deficient, lineage bias could be determined upstream of the oligopotent progenitors.

**Figure 6.**
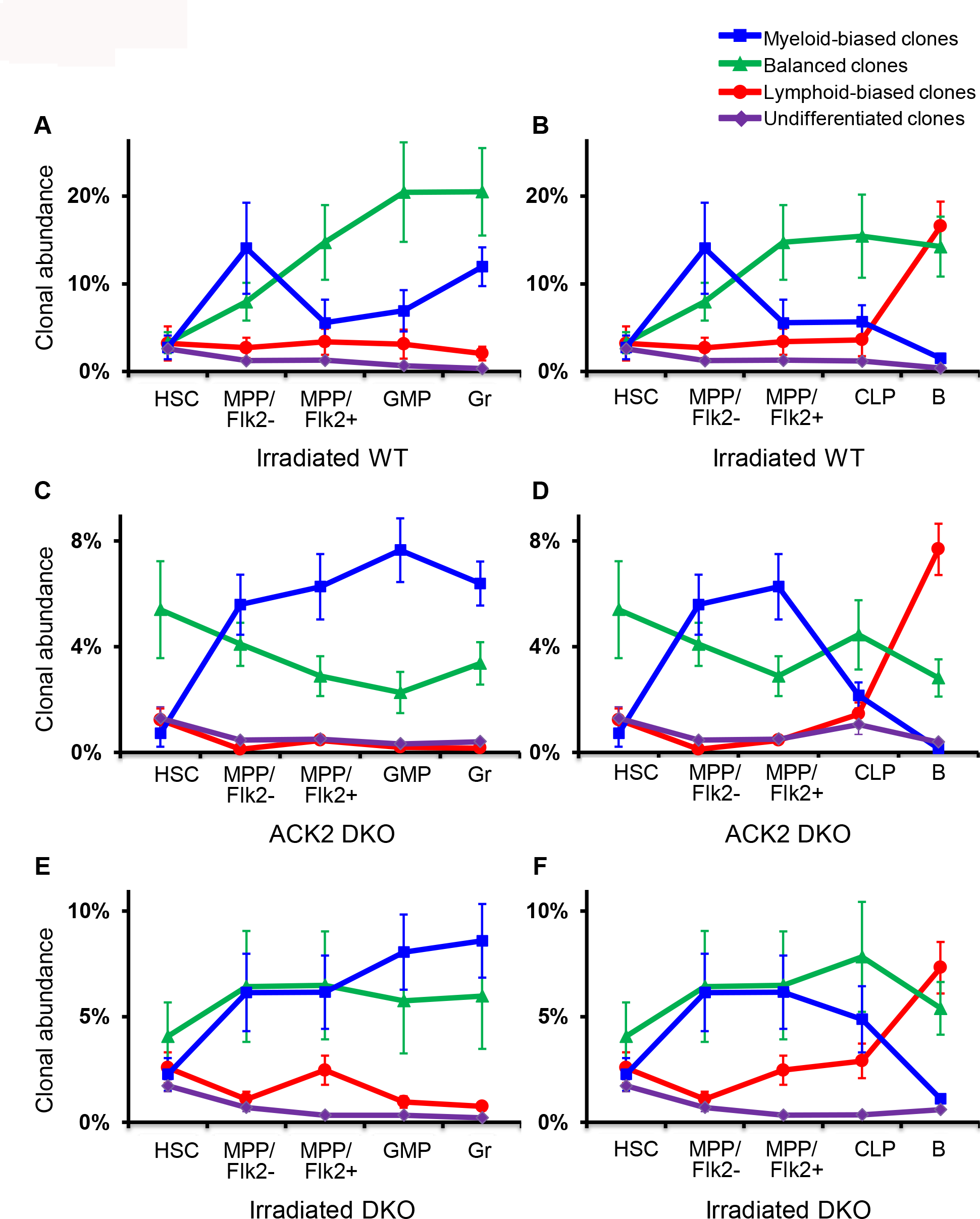
Lineage Commitment of HSC Clones With Distinct Lineage Bias and Lineage Balance. Shown are the average abundance of clones with distinct lineage bias and balance at various stages of HSC differentiation. Clonal abundance here refers to the copy number of a particular barcode as a percentage of all barcode copy numbers from a cell population. (A-B) Irradiated wild type mice. (C-D) ACK2-treated Rag2^−/−^ϒc^−/−^ mice. (E-F) Irradiated Rag2^−/−^ϒc^−/−^ mice. (A, C and E) Myeloid differentiation towards granulocytes. (B, D and F) Lymphoid differentiation towards B cells. Error bars show the standard errors of the means for all the barcodes from mice under the same transplantation conditions. WT: wild type mice. DKO: Rag2^−/−^ϒc^−/−^ mice.

### Lineage Commitment Profiles of HSC Clones With Distinct Lineage Bias and Balance

To systematically examine the lineage commitment paths that lead to differential blood production, we analyzed the average abundance of clones with distinct lineage bias and balance through various stages of lineage commitment (Fig. 6). In ACK2-treated DKO mice, myeloid-biased clones expand dominantly between the HSC and MPP^Flk2−^ stages (Fig. 5A and 6C). These clones are diminished at the CLP and B cell stages (Fig. 6D). In contrast, lymphoid-biased clones expand dominantly between the CLP and B cell stages (Fig. 5A and 6D), and their abundances remain constant during the initial lineage commitment and the myeloid lineage commitment stages (Fig. 6C). Balanced HSC clones do not exhibit dominant differentiation in either lineage (Fig. 6C–D), consistent with previous analysis (Fig. 4B and S5). The fluctuations of the balanced clones appear to be inversely correlated with the dominant differentiation of myeloid-biased clones and lymphoid-biased clones (Fig. 6C–D), indicating that the fluctuation is not likely to be a real change but is rather a reflection of the dominant expansion of other clones. This can be attributed to the data normalization procedure that normalizes the total abundance of all barcodes in each cell population to 100%. Interestingly, these balanced clones are more abundant at the HSC stage than other clones (Fig. 4E and 6C–D). The high abundances persist in their downstream progenies (Fig. 6C–D), suggesting that balanced clones are significantly more committed to self-renewal than other clones, and that all self-renewed balanced HSCs differentiate.

In irradiated wild type mice, lineage-biased and balanced clones all exhibit similar abundances at the HSC stage (Fig. 6A–B). Myeloid-biased clones dominantly expand twice during the lineage commitment process (Fig. 6A). The first dominant expansion occurs between the HSC and MPP^Flk2−^ stages and the second dominant expansion occurs between the GMP and granulocyte stages (Fig. 6A). There is a retrenchment between these two expansions, and irradiated wild type mice do not exhibit lineage bias at the progenitor stages (Fig. 5D). Myeloid-biased clones do not expand between the CLP and B cell stages, whereas lymphoid-biased clones expand dominantly at this lineage commitment step (Fig. 5A and 6B). Lymphoid-biased clones do not dominantly expand at any other lineage commitment steps, but they are more abundant than undifferentiated clones at the MPP and CLP stages, suggesting that lymphoid-biased clones are not absent during the early stages of lymphoid differentiation (Fig. 6A–B). Most strikingly, balanced clones dramatically expand in irradiated mice at almost every lineage commitment step (Fig. 6A–B). Their relatively less severe expansion at the last step, the GMP-to-granulocyte step and the CLP-to-B-cell step, is likely a reflection of the dominant differentiation of myeloid-biased clones and lymphoid-biased clones, due to data normalization (Fig. 6A–B).

In irradiated DKO mice, balanced clones exhibit moderate expansion during self-renewal and differentiation (Fig. 6E–F), a phenotype in between that of ACK2-treated DKO mice and irradiated wild-type mice (Fig. 6A–D). Myeloid-biased clones and lymphoid-biased clones in irradiated DKO mice (Fig. 6E–F) exhibit similar lineage commitment profiles as those in ACK2-treated DKO mice (Fig. 6C–D), combining characteristics from irradiated wild type mice (Fig. 6A–B) such as the double expansion of myeloid-biased clones. The second expansion of myeloid-biased clones occurs not at the last lineage commitment step, but at the MPP^Flk2+^-to-GMP step. In addition, lymphoid-biased clones also start to expand early at the MPP^Flk2−^-to-MPP^Flk2+^ step (Fig. 6E–F). The early expansion of lineage-biased clones in irradiated DKO mice may be related to the ubiquitous expansion of all differentiating clones observed in irradiated wild type mice. Taken together, the comparison of irradiated DKO mice and ACK2-treated DKO mice manifests the characteristics of irradiation-mediated transplantation.

### A Model of Clonal Level Lineage Commitment Pathways of HSCs *In Vivo*

Based on all of the data above (Fig. 1–6), we propose a model for the clonal level lineage commitment pathways of HSCs *in vivo* (Fig. 7). After unconditioned transplantation, all engrafted HSCs uniformly differentiate and self-renew (Fig. 1B–C, 2A–B and 2G). While they may contribute differently to blood cells with distinct maturation processes, such as T cells (Fig. 1D–E), they do not exhibit dominant differentiation nor myeloid versus lymphoid lineage bias (Fig. 2–3). In contrast, after conditioned transplantation, only a small subset of engrafted HSC clones is involved in differentiation (Fig. 2C–F). These clones follow distinct pathways that are characterized by dominant differentiation and lineage bias (Fig. 7).

**Figure 7.**
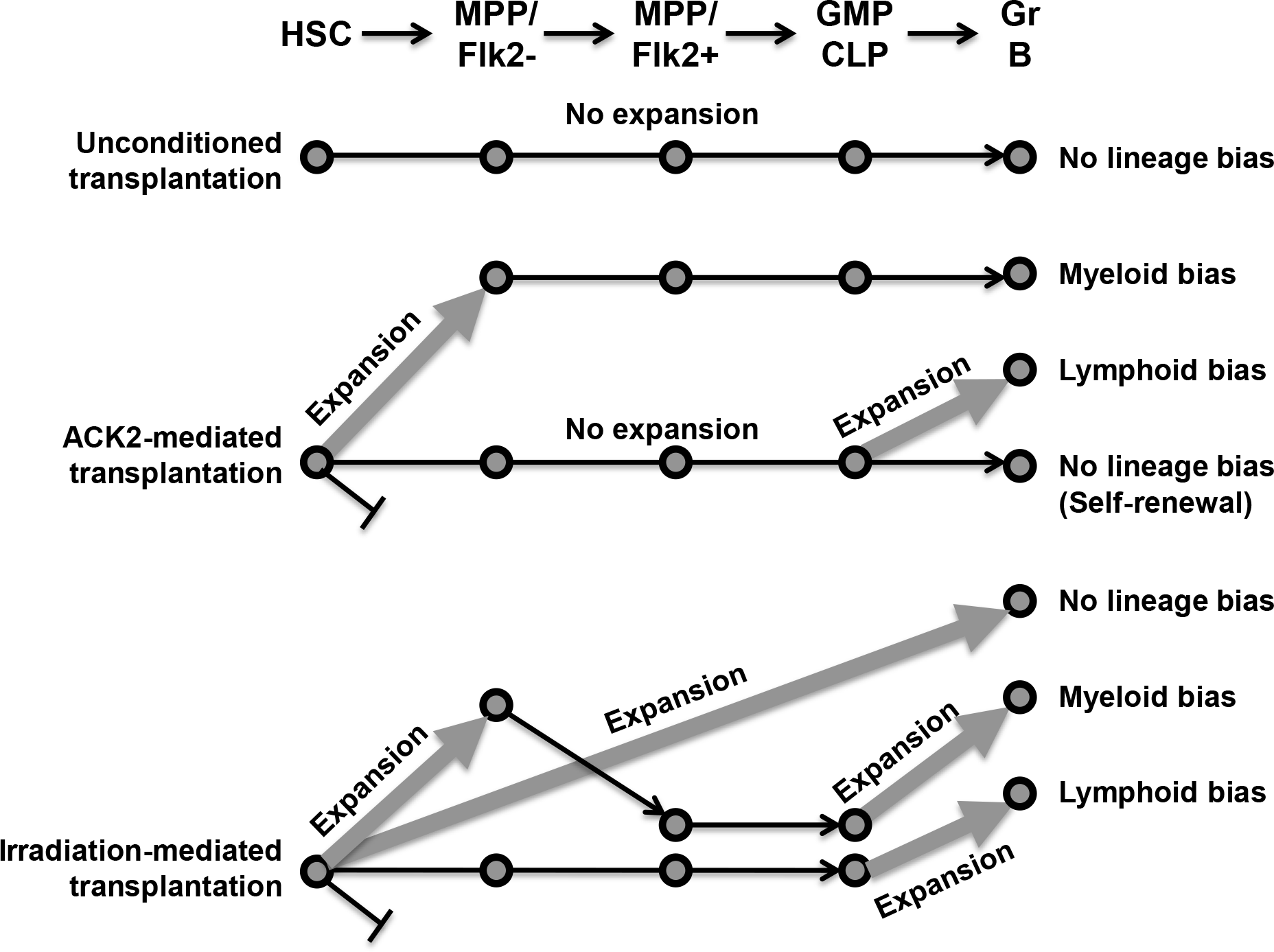
A Model of Clonal Level Lineage Commitment Pathways of HSCs *In Vivo*. Thicker and ascendant arrows represent dominant differentiation of HSC clones. HSCs transplanted into mice pre-treated with different conditioning regimens follow distinct pathways during lineage commitment. These pathways lead to distinct balanced or biased blood production.

After ACK2 antibody mediated transplantation, differentiating HSC clones follow one of three pathways (Fig. 7). In the first pathway, HSC clones do not exhibit dominant differentiation nor lineage bias (Fig. 4B). Their lineage commitment process resembles that of HSC clones transplanted into unconditioned recipients (Fig. 4D). Furthermore, these HSC clones self-renew significantly more than other HSC clones (Fig. 4E and 6C–D). In the second pathway, HSC clones dominantly expand at the first lineage commitment step, HSC to MPP^Flk2−^ (Fig. 5A). These HSC clones eventually become myeloid-biased (Fig. 5B). In the third pathway, HSC clones dominantly expand at the last lineage commitment step, CLP to B cell (Fig. 5A) and end up with lymphoid bias (Fig. 5B). A significant number of HSC clones are not recruited into differentiation and do not participate in any of these pathways (Fig. 6C–D and S6E–F).

After irradiation-mediated transplantation, HSC clones follow similar lineage commitment pathways as those after ACK2-mediated transplantation (Fig. 7). The major difference is that all differentiating HSCs dominantly expand during differentiation in irradiated recipients regardless of their lineage bias or balance (Fig. 6A–B). This dominant expansion is manifested in distinct differentiation lineages and stages in different pathways. However, none of these pathways are associated with dominant self-renewal (Fig. 4E and 6A–B). In particular, balanced HSCs expand dramatically at every step of the lineage commitment process (Fig. 6A–B). The dominant differentiation of balanced clones may restrain the expansion of myeloid-biased clones, such that myeloid-biased clones are not significantly more present in GMP than lymphoid-biased clones (Fig. 6A). Therefore, lineage bias is absent at the GMP and CLP stages in wild-type irradiated recipients (Fig. 5D). Downstream of these progenitors, myeloid-biased clones expand again at the last step of lineage commitment from GMP to granulocyte. Lymphoid-biased and undifferentiated pathways in irradiated recipients exhibit similar lineage commitment characteristics as their counterparts in ACK2-treated recipients (Fig. 5A–B and 6).

## DISCUSSION

In this study, we identified the clonal level lineage commitment pathways of hematopoietic stem cells *in vivo* (Fig. 7). We show that lineage commitment of HSC clones after conditioning is characterized by dominant differentiation and lineage bias (Fig. 2–3). Dominant differentiation and lineage bias are interrelated (Fig. 4–5) and together delineate distinct pathways that lead to balanced or biased blood production (Fig. 6–7). These pathways elucidate cellular proliferation and development of HSCs at the clonal level and demonstrate distinct modes of HSC regulation *in vivo*.

### Pre-transplantation Conditioning Alters HSC Differentiation at the Clonal Level

Irradiation is used in virtually all HSC studies. It is also widely applied in clinical therapy to facilitate bone marrow transplantation and to treat cancers and hematopoietic disorders. Here, we have shown how irradiation alters HSC regulation at the clonal level (Fig. 2–3). This striking alteration could lead to new interpretations of studies of HSC physiology that use irradiation as a conditioning regimen. For example, many recent studies propose that HSCs are heterogeneous and possess differential lineage bias (8, 10–15). These studies all use irradiation to facilitate HSC engraftment. Our data now demonstrate that engrafted HSCs uniformly differentiate and self-renew in the absence of any pre-transplantation conditioning and that heterogeneous hematopoiesis is only observed after conditioned transplantation (Fig. 2–3). This indicates that the conditioning regimen used in the previous studies may have contributed to the observed HSC heterogeneity. Thus, future studies must be carefully designed to distinguish normal HSC physiology from emergency modes.

HSC regulatory mechanisms activated after conditioning are likely to be more susceptible to perturbation and damage (43). These mechanisms may be key to understanding how hematopoiesis becomes malignant and for reducing the side effects of clinical regimens used to treat these malignancies. For example, several gene therapy trials were dismayed by the appearance of clonal dominance in the blood cells of treated patients (44, 45). This clonal dominance was interpreted to be a result of viral integration that ectopically activates nearby oncogenes and drives cellular expansion. However, our data suggests that the observed clonal dominance may instead have been induced by the use of pre-transplantation conditioning regimens that accompanied the gene therapy procedure. With this new knowledge in mind, gene therapists may wish to try various conditioning regimens in order to achieve the optimal clinical outcome.

### Cellular Origins of Clonal Dominance and Lineage Bias

Our elucidation of the HSC clonal level lineage commitment pathways reveals the new HSC regulatory mechanisms and identifies the cellular origins of clonal dominance and lineage bias that were previously observed in blood cells (8, 10–13, 15, 46–48). We show that clonal dominance is initiated by the outgrowth of clones predominantly at the hematopoietic progenitor stages, and not at the HSC stage (Fig. 2). This explains why previous studies had detected few clones in the blood post irradiation (11, 46–48). We demonstrate that lineage bias arises from the dominant clonal expansion of specific lineages at key lineage commitment steps in conditioned mice (Fig. 5) and explains the presence of lineage bias in blood cells (8, 10, 12, 13, 15).

### Unconditioned Transplantation Provides Unique Insights into Natural HSC Physiology

Unconditioned transplantation minimally perturbs natural hematopoiesis and provides new insights into natural HSC physiology. We did not detect the presence of “dormant HSCs” (49) in unconditioned transplantation (Fig. 1B–C and 2A–B). It is possible that “active HSCs” are selectively engrafted or that HSCs engrafted after unconditioned transplantation are regulated as active HSCs and not as dormant HSCs. Nonetheless, our data suggest that engrafted HSCs continuously and homogenously contribute to the blood pool (Fig. 1B–C and 2A–B). As HSCs frequently migrate through the peripheral blood and re-engraft under natural homeostatic conditions (33–35), this process may be a natural procedure to select for active and lineage-balanced HSCs (Fig. 1B–C and 2A–B). The homogeneity of HSC clonal behavior indicates that HSCs can be uniformly regulated and do not require a complex regulatory system to shepherd selected HSCs in and out of differentiation and self-renewal under homeostatic conditions (11, 47, 48, 50).

### Different Clonal Pathways Can Coexist Simultaneously in a Single Organism

By tracking many HSC clones simultaneously, we identified several distinct pathways co-existing in a single mouse after conditioning (Fig. 7). These pathways mutually compensate to sustain overall blood production (Fig. 6–7). Thus, the striking differences in HSC regulation uncovered by this clonal analysis are not evident at the population level. This is not unexpected, as blood cells are critical for the survival of the organism. Redundant and feedback mechanisms have evolved to maintain overall blood production (2, 51).

An unexpected co-existence of different clonal pathways is found in ACK2-treated mice (Fig. 4B, 4D and 7). In these mice, one pathway preserves the characteristics from the unconditioned state, lacking both dominant differentiation and lineage bias. The other pathways resemble those from the irradiated state, exhibiting both dominant differentiation and linage bias. The presence of the unconditioned pathway may be responsible for the long-term benefit of the non-myeloablative transplantation observed in clinic therapy (52, 53), as irradiation and ACK2 treatments produce similar degrees of dominant differentiation and lineage bias (Fig. 2–3 and S3C). The co-existence of the unconditioned pathway with pathways activated after conditioning in the same mouse suggests that these pathways are not mutually exclusive and are not altered by global mobilized factors. It also provides the possibility for activating the unconditioned pathway in myeloablated (irradiated) patients as part of a therapeutic procedure to achieve similar benefits to that of non-myeloablated patients.

### Molecular Mechanisms Underlying the Lineage Commitment Pathways of HSC Clones

The molecular mechanisms underlying the lineage commitment pathways of HSC clones are complex (Fig. 7). They likely involve both intrinsic and extrinsic factors. Several recent studies have revealed cell surface markers on HSCs that are enriched with distinct lineage bias in irradiated hosts (8, 12, 15, 54, 55). This suggests that select HSCs preferentially follow certain pathways. This outcome is complex, as the myeloid-biased HSC clones are enriched with CD150^hi^, and the balanced clones are enriched with CD150^med^ (8, 12, 15, 55), showing that the predisposition for clonotypes is determined at the HSC level, but read out at the level of different progenitors for the inclusion or relative exclusion of CLP and progeny in the mature cell pool. It has also been shown that cytokines can direct hematopoiesis into distinct lineages (4–7), which suggests that extrinsic factors can alter the pathway choices of HSCs.

HSCs in conditioned recipients receive different regulatory signals compared with those in unconditioned recipients (Fig. 2–3). These signals may either selectively engraft a subset of HSCs that are lineage-biased or may induce lineage bias from balanced HSCs. In the latter scenario, it is possible that intrinsic differences of HSCs elicit different responses to the conditioning regimen (8, 12, 15, 54, 55). These regulatory signals may be produced as a consequence of stimulation or damage to HSC niche by the conditioning regimen (27). A deficiency of blood cells may also induce feedback signals that boost HSC differentiation. These signals can activate the first few HSCs landing at niches and instruct them to dominantly differentiate to compensate for the blood loss. We have shown that after irradiation all differentiating HSC clones expand dominantly at various stages (Fig. 6A–B and 7), indicating the stress for the HSCs to supply the blood cells after irradiation.

In summary, we have provided a comprehensive view of HSC lineage commitment at the clonal level for the first time and have uncovered the underlying lineage commitment pathways of individual HSC clones (Fig. 7). These pathways are altered by transplantation conditioning such as irradiation (Fig. 2–3), which has been ubiquitously used in HSC studies. The pathways also identify the HSC differentiation stages where HSC clones become dominant and where lineage bias originates (Fig. 5–6). Knowledge of clonal level HSC lineage commitment pathways opens new avenues of research for understanding and manipulating blood production and balance.

## EXPERIMENTAL PROCEDURES

The donor mice used in all experiments were C57BL6/Ka (CD45.1^+^). The recipient mice used in the unconditioned transplantation experiments (M1-8) were C57BL6/Ka (CD45.1^+^/ CD45.2^+^). The recipient mice used in the irradiation-mediated transplantation experiments (M9-15, and those used in Fig. 3E) were C57B L6/Ka (CD45.2^+^). The recipient mice used in the ACK2-mediated transplantation experiments (M16-22) were Rag2^−/−^ϒc^−/−^ with C57BL6/Ka background (DKO) (26). The DKO mice were also used as irradiation recipients (M23-32). All donor and recipient mice were 8-12 weeks old at the time of transplantation. Mice were bred and maintained at Stanford University’s Research Animal Facility. Animal procedures were approved by the International Animal Care and Use Committee.

HSCs (lineage (CD3, CD4, CD8, B220, Gr1, Mac1, Ter119)^−^/ckit^+^/Sca1^+^/Flk2^−^/CD34^−^/CD150^+^) were obtained from the crushed bones of donor mice and isolated using double FACS sorting with the FACS-Aria II (BD Biosciences, San Jose, CA) after enrichment using CD117 microbeads (AutoMACS, Miltenyi Biotec, Auburn, CA). HSCs were infected for 10 hours with lentivirus carrying barcodes and then transplanted via retro-orbital injection. Recipient mice were treated with one of the following three conditions before transplantation: (a) no treatment, referred to as “unconditioned” (M1-8); (b) irradiation with 950 cGy immediately prior to transplantation (M9-15 and M23-32); (c) retro-orbital injection of 500 µg of ACK2 into Rag2^−/−^ϒc^−/−^ mice 9 days prior to transplantation (26) (M16-22). In unconditioned transplantation, 1000 barcoded HSCs were transplanted into each mouse every other day for eighteen days (9000 donor HSCs total). Long-term stable engraftment of approximately 1% donor chimerism was consistently obtained. In irradiation-mediated and ACK2-mediated transplantation, 9000 donor HSCs were transplanted all at once. Donor chimerisms obtained were around 10% and 90% respectively. 250,000 whole bone marrow cells without viral transduction were co-transplanted into each irradiated mouse as helper cells, except specified otherwise (Fig. 3E). All cells were harvested 22 weeks after transplantation. HSC clonal labeling and data analysis are explained in detail elsewhere (36). Dominant clones are defined as those whose relative copy numbers in blood cells are more than five times their relative copy numbers in HSCs. Lineage-biased clones are defined as those whose relative copy numbers in one lineage are more than 2.4142 (cotangent 22.5 degree) times their relative copy numbers in the other lineage. Low-abundance clones are excluded from the analysis of lineage bias versus balance. These clones are defined as those whose copy numbers are less than 10% of the maximum copy numbers in both lineages. All results have been reaffirmed using different lineage bias and clonal dominance threshold values.

Below is the list of cell surface markers used to harvest hematopoietic populations. Donor cells were sorted based on the CD45 marker.

Granulocytes: CD4^−^/CD8^−^/B220^−^/CD19^−^/Mac1^+^/Gr1^+^/side scatter^high^;

B cells: CD4^−^/CD8^−^/Gr1^−^/Mac1^−^/B220^+^/CD19^+^;

CD4 T cells: B220^−^/CD19^−^/Mac1^−^/Gr1^−^/TCRαβ^+^/CD4^+^/CD8^−^;

CD8 T cells: B220^−^/CD19^−^/Mac1^−^/Gr1^−^/TCRαβ^+^/CD4^−^/CD8^+^;

HSC (hematopoietic stem cells): lineage (CD3, CD4, CD8, B220, Gr1, Mac1, Ter119)^−^/IL7Rα^−^/ckit^+^/Sca1^+^/Flk2^−^/CD34^−^/CD150^+^;

MPP^Flk2−^ (Flk2−multipotent progenitor): lineage (CD3, CD4, CD8, B220, Gr1, Mac1, Ter119)^−^/IL7Rα^−^/ckit^+^/Sca1^+^/Flk2^−^/CD34+;

MPP^Flk2+^ (Flk2+ multipotent progenitor): lineage (CD3, CD4, CD8, B220, Gr1, Mac1, Ter119)^−^/IL7Rα^−^/ckit^+^/Sca1^+^/Flk2^+^;

CLP (common lymphocyte progenitor): lineage (CD3, CD4, CD8, B220, Gr1, Mac1, Ter119)^−^/IL7Rα^+^/Flk2^+^;

GMP (granulocyte/monocytic progenitor): lineage (CD3, CD4, CD8, B220, Gr1, Mac1, Ter119)^−^/IL7Rα^−^/ckit^+^/Sca1^−^/CD34^+^/FcϒR^+^.

## ACKNOWLEDGEMENTS

We thank N. Neff, G. Mantalas, T. Snyder, B. Passarelli and S. Quake for carrying out the high-throughput sequencing; thank M. Inlay, T. Serwold and C. Chan for helpful discussions on experiments; and thank H. Nakauchi, S. Karten, K. Loh for helpful suggestions on the manuscript. We also thank L. Jerabek and T. Storm for laboratory management; C. Muscat and T. Naik for antibody conjugation; A. Mosley for animal supervision; P. Lovelace for FACS core management and technical support. This work is supported by NIH-R01-CA86065, NIH-U01-HL099999. R.L. is supported NIH K99-HL113104, NIH R00-HL113104, R01HL135292, R01HL138225 and P30CA014089.

R.L., A.C., J.S. and I.L.W. designed the experiments. R.L. and D.J. performed the experiments and analyzed the data. R.L. wrote the manuscript. All authors edited the manuscript.

---

**Supplemental Figure 1.**
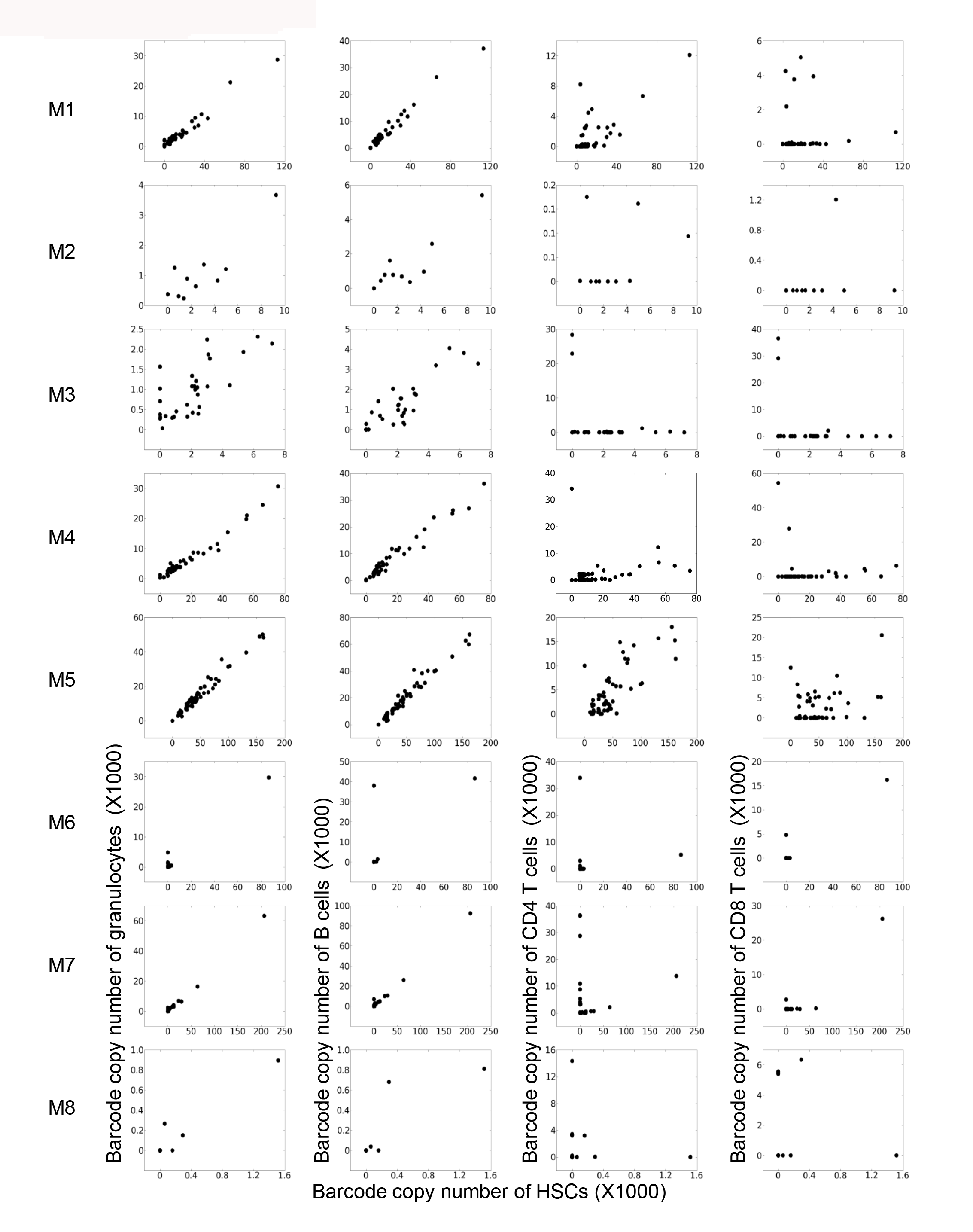
Comparing Clonality of HSCs with Clonality of Blood Cells in Individual Mice. Related to Figure 1B–E. Plots were generated as described in Figure 1 B–E’s legend. These scatter plots compare the copy numbers of barcodes from HSCs (x-axis) with barcodes from different blood cell types (y-axis). Each plot depicts a single mouse twenty-two weeks after unconditioned transplantation. The plots illustrate the linear correlations of barcode copy numbers from HSCs with those from granulocytes and B cells. Note that some HSC barcodes do not appear in CD4 T cells nor in CD8 T cells from the peripheral blood.

**Supplemental Figure 2.**
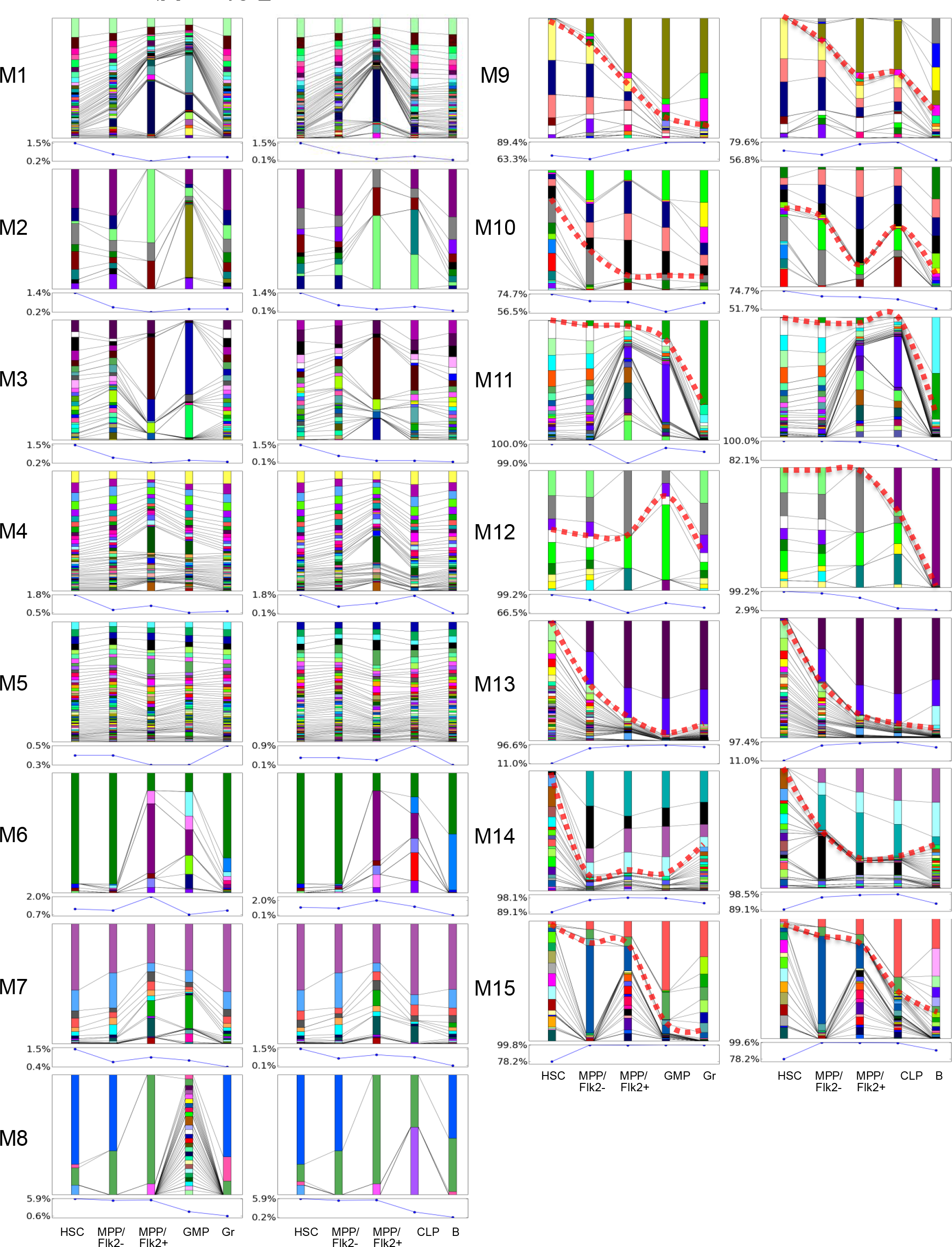
Dominant Expansion During Myeloid Differentiation and Lymphoid Differentiation in Individual Mice. Related to Figure 2A–D. Plots were generated as described in Figure 2 A–D’s legend. Each plot illustrates HSC lineage commitment within a single mouse. The line plot under each column plot depicts the donor chimerism measured using the CD45 marker. (M1-8) unconditioned mice; (M9-15) lethally irradiated mice. The data suggests that clonal dominance occurs during myeloid differentiation and lymphoid differentiation after irradiation-mediated transplantation but not after unconditioned transplantation. The clonal expansion profile is correlated with the donor chimerism profile.

**Supplemental Figure 3.**
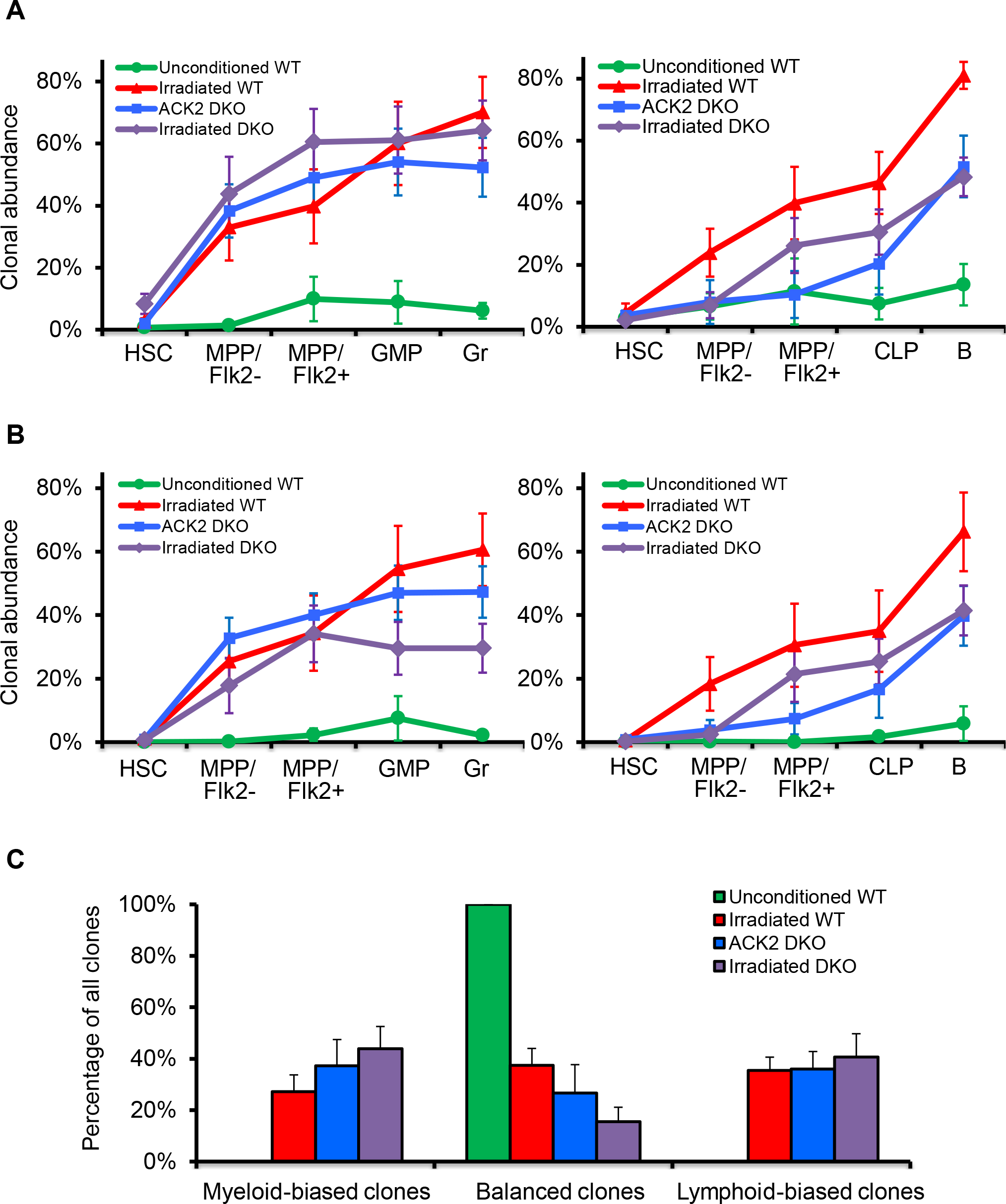
Dominant Differentiation and Lineage Bias in Mice Treated With Different Transplantation Conditioning. Related to Figure 2E–F and 3A–D. (A-B) Plots were generated as described in Figure 2E–F’s legend, except that the threshold value defining dominant clones is different. (A) Dominant clones are defined as clones whose relative copy numbers in the peripheral blood cells are more than twice their relative copy numbers in HSCs. (B) Dominant clones are defined as clones whose relative copy numbers in the peripheral blood cells are more than ten times their relative copy numbers in HSCs. (C) Percentage of HSC clones that are lineage-biased or lineage-balanced under different conditioning regimen. WT: wild type mice. DKO: Rag2^−/−^ϒc^−/−^ mice. (A-C) Error bars represent the standard errors of the means of all mice under identical transplantation conditions. The data show that HSCs exhibit dominant differentiation and lineage bias after conditioned transplantation but not after unconditioned transplantation.

**Supplemental Figure 4.**
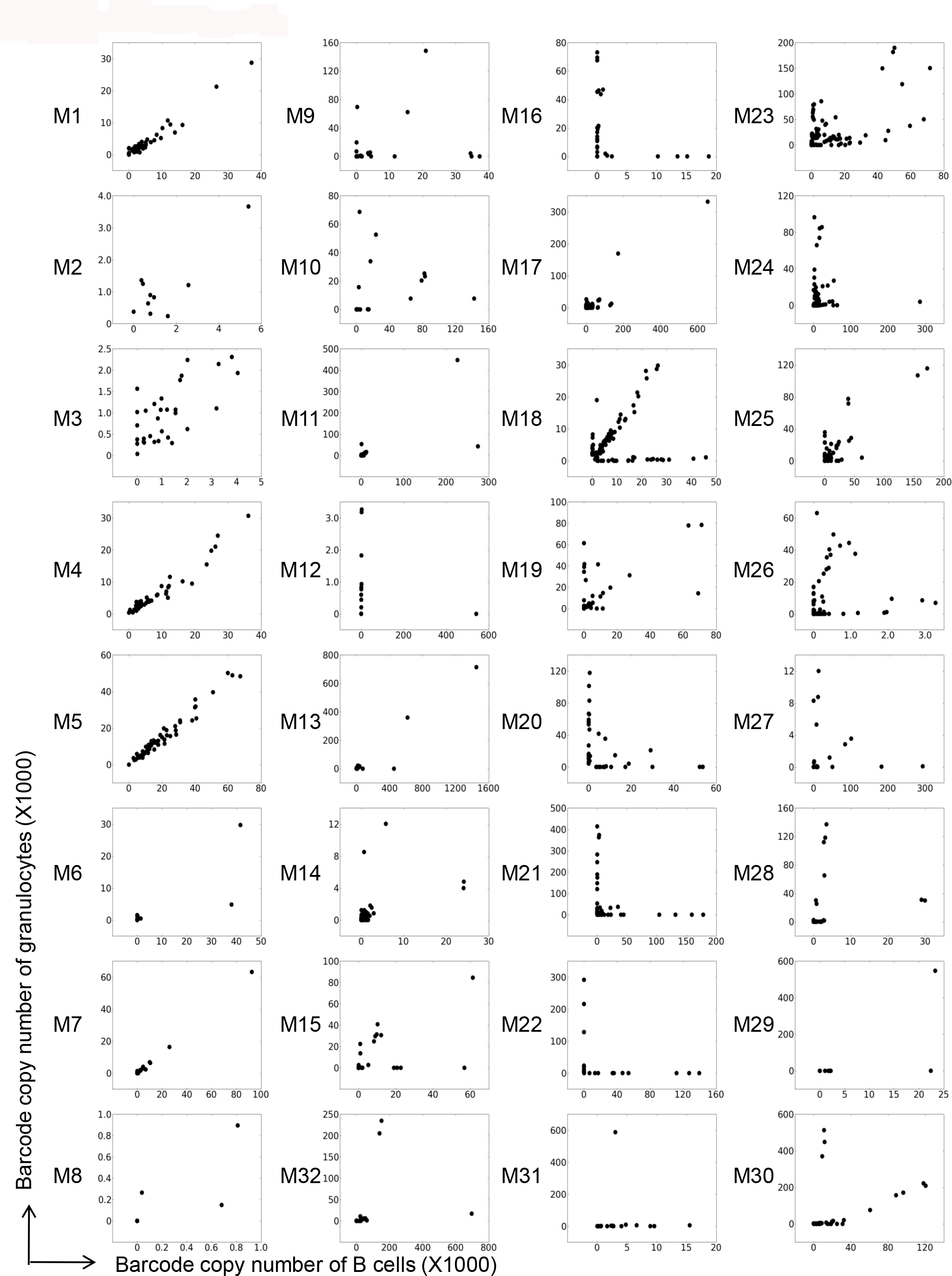
Comparing Clonality of Granulocytes with Clonality of B Cells in Individual Mice. Related to Figure 3A–D. Plots were generated as described in Figure 1 B–E’s legend. These scatter plots compare the copy numbers of barcodes from granulocytes (y-axis) with barcodes from B cells (x-axis). Each plot depicts a single mouse twenty-two weeks after transplantation. (M1-8) wild type mice transplanted without any conditioning; (M9-15) wild type mice transplanted after lethal irradiation; (M16-22) Rag2^−/−^ϒc^−/−^ mice transplanted after ACK2 treatment; (M23-32) Rag2^−/−^ϒc^−/−^ mice transplanted after lethal irradiation. The data suggests that HSCs exhibit lineage bias after conditioned transplantation but not after unconditioned transplantation.

**Supplemental Figure 5.**
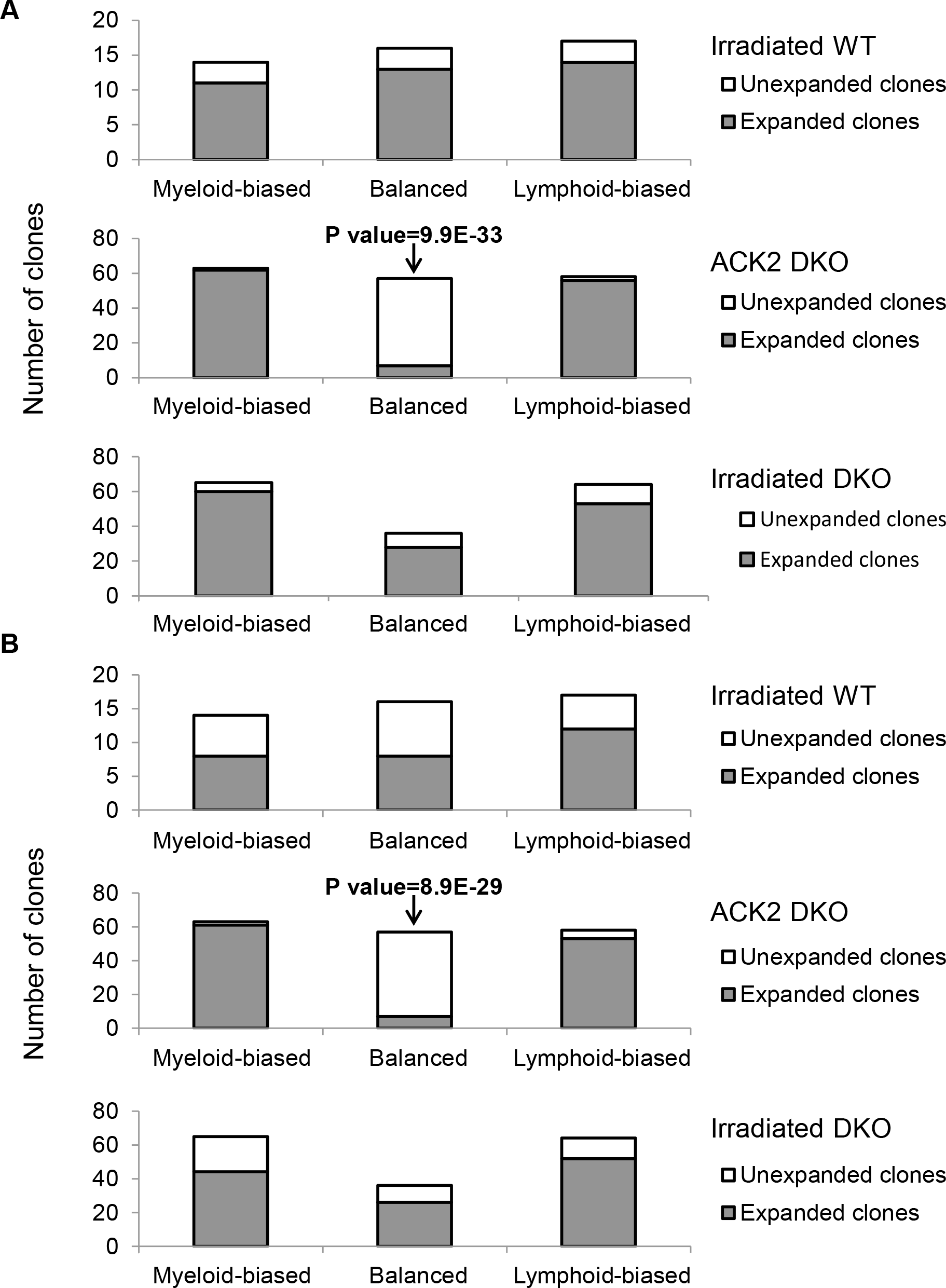
Lineage Bias and Balance of Dominant Clones and Non-dominant Clones. Related to Figure 4A–C. (A-B) Plots were generated as described in Figure 4A–C’s legend, except that the threshold value defining dominant clones is different. (A) Dominant clones are defined as the clones whose relative copy numbers in peripheral blood cells are more than twice their relative copy numbers in HSCs. (B) Dominant clones are defined as clones whose relative copy numbers in the peripheral blood cells are more than ten times their relative copy numbers in HSCs. The data suggests that dominant clones exhibit lineage bias and non-dominant clones exhibit lineage balance in ACK2 treated mice. However, this correlation is not observed in irradiated mice. *** P<0.001.

**Supplemental Figure 6.**
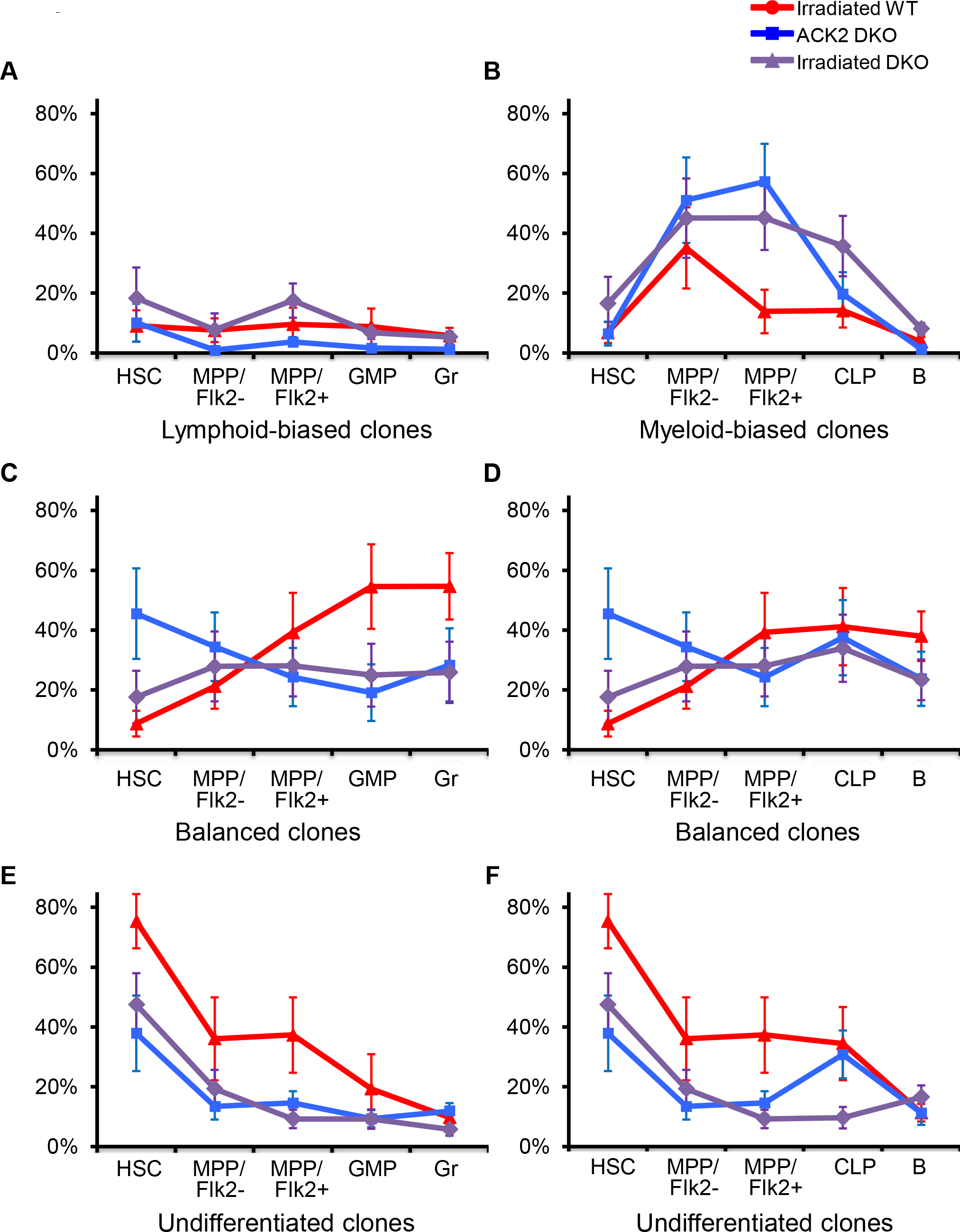
Lineage Commitment of Clones with Distinct Lineage Bias and Balance. Related to Figure 5A–B. Plots were generated as described in Figure 5A–B’s legend. Shown is the percentage of barcodes representing distinct lineage bias and balance at various stages of differentiation. Error bars represent the standard errors of the means of all mice under identical transplantation conditions. The data suggests that lymphoid-biased clones remain unexpanded in myeloid differentiation (A) and myeloid-biased clones diminish in lymphoid lineage (B). Balanced clones stably differentiate after ACK2 treatment (C and D). But they dominantly differentiate in irradiated wild type mice (C and D). Undifferentiated clones form a large portion of the HSC population (E and F). Their abundances decrease in the differentiated populations (E and F).

**Supplemental Figure 7.**
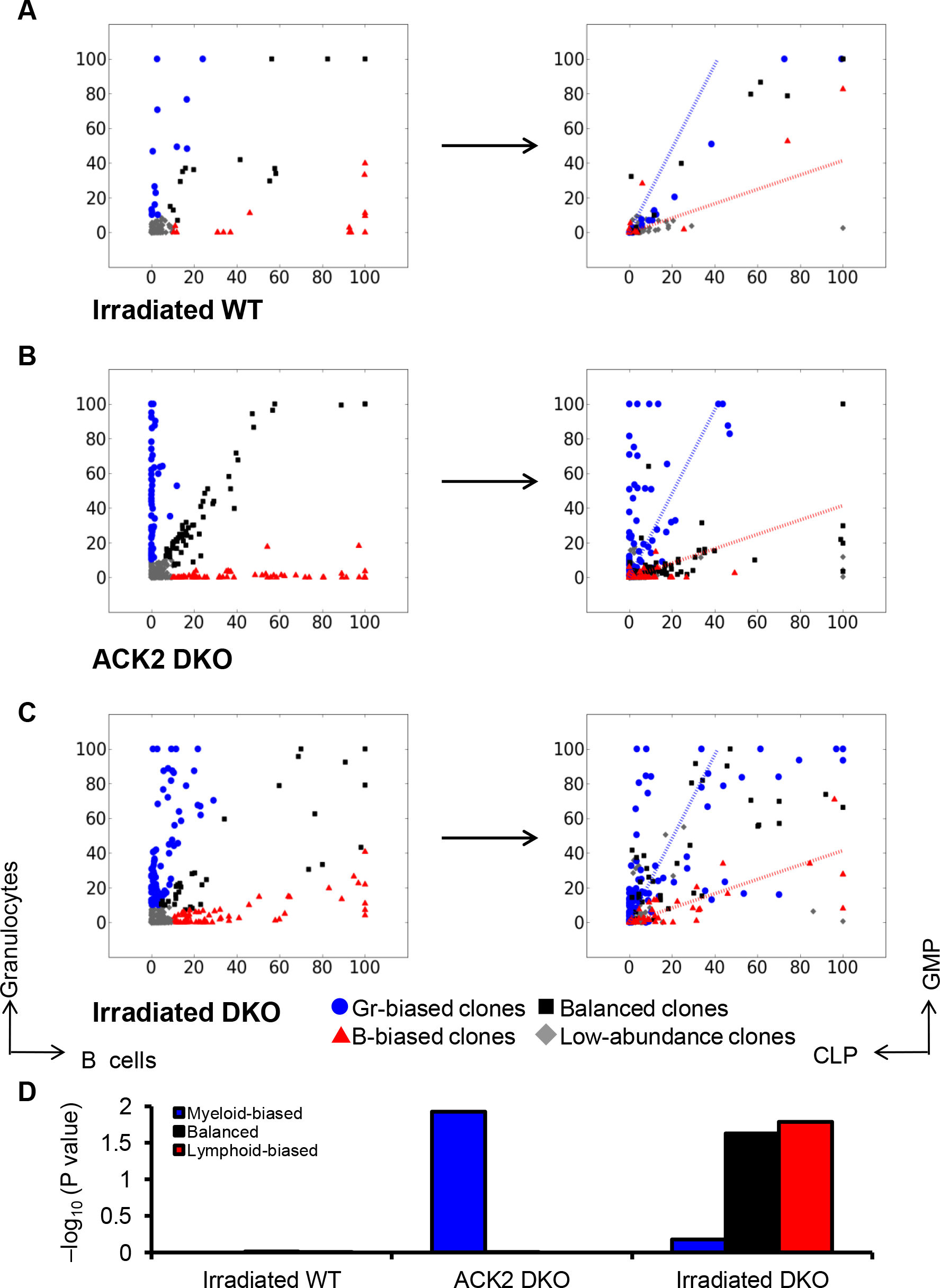
Comparing Lineage Bias of Mature Blood Cells with Lineage Bias of Progenitors. Related to Figure 5C–D. Plots were generated as described in Figure 5C–D’s legend, except that dot colors are assigned based on the bias between granulocytes and B cells. (A-C right) The dotted lines show the boundary of lineage bias and balance in progenitor cells. (D) P value depicts the significance that the lineage bias and balance in the blood cells is determined at the progenitor level. P value is calculated based on the assumption that the clones are randomly distributed among the different categories of lineage bias and balance. The data suggests that the lineage bias of mature blood cells is only correlated with the lineage bias of progenitors in Rag2^−/−^ϒc^−/−^ recipients. This correlation does not exist in in wild type recipients.

## References

1. Bryder D, Rossi DJ, Weissman IL (2006) Hematopoietic stem cells: the paradigmatic tissue-specific stem cell. Am J Pathol 169(2):338–346.

2. Seita J, Weissman IL (2010) Hematopoietic stem cell: self-renewal versus differentiation. Wiley Interdiscip Rev Syst Biol Med 2(6):640–653.

3. Weissman IL (2000) Stem cells: units of development, units of regeneration, and units in evolution. cell 100(1):157–168.

4. Kondo M, et al. (2000) Cell-fate conversion of lymphoid-committed progenitors by instructive actions of cytokines. Nature 407(6802):383–386.

5. Rankin EB, et al. (2012) The HIF signaling pathway in osteoblasts directly modulates erythropoiesis through the production of EPO. Cell 149(1):63–74.

6. Rieger MA, Hoppe PS, Smejkal BM, Eitelhuber AC, Schroeder T (2009) Hematopoietic cytokines can instruct lineage choice. Science 325(5937):217–218.

7. Sarrazin S, et al. (2009) MafB restricts M-CSF-dependent myeloid commitment divisions of hematopoietic stem cells. Cell 138(2):300–313.

8. Beerman I, et al. (2010) Functionally distinct hematopoietic stem cells modulate hematopoietic lineage potential during aging by a mechanism of clonal expansion. Proc Natl Acad Sci 107(12):5465–5470.

9. Copley MR, Beer PA, Eaves CJ (2012) Hematopoietic stem cell heterogeneity takes center stage. Cell Stem Cell 10(6):690–697.

10. Dykstra B, et al. (2007) Long-term propagation of distinct hematopoietic differentiation programs in vivo. Cell Stem Cell 1(2):218–229.

11. McKenzie JL, Gan OI, Doedens M, Wang JC, Dick JE (2006) Individual stem cells with highly variable proliferation and self-renewal properties comprise the human hematopoietic stem cell compartment. Nat Immunol 7(11):1225–1233.

12. Morita Y, Ema H, Nakauchi H (2010) Heterogeneity and hierarchy within the most primitive hematopoietic stem cell compartment. J Exp Med 207(6):1173–1182.

13. Muller-Sieburg CE, Cho RH, Karlsson L, Huang J-F, Sieburg HB (2004) Myeloid-biased hematopoietic stem cells have extensive self-renewal capacity but generate diminished lymphoid progeny with impaired IL-7 responsiveness. Blood 103(11):4111–4118.

14. Sieburg HB, et al. (2006) The hematopoietic stem compartment consists of a limited number of discrete stem cell subsets. Blood 107(6):2311–2316.

15. Weksberg DC, Chambers SM, Boles NC, Goodell MA (2008) CD150-side population cells represent a functionally distinct population of long-term hematopoietic stem cells. Blood 111(4):2444–2451.

16. Sun J, et al. (2014) Clonal dynamics of native haematopoiesis. Nature 514(7522):322–327.

17. Lu R (2014) Sleeping beauty wakes up the clonal succession model for homeostatic hematopoiesis. Cell Stem Cell 15(6):677–678.

18. Rodriguez-Fraticelli AE, et al. (2018) Clonal analysis of lineage fate in native haematopoiesis. Nature.

19. Yamamoto R, et al. (2013) Clonal analysis unveils self-renewing lineage-restricted progenitors generated directly from hematopoietic stem cells. Cell 154(5):1112–1126.

20. Cho RH, Sieburg HB, Muller-Sieburg CE (2008) A new mechanism for the aging of hematopoietic stem cells: aging changes the clonal composition of the stem cell compartment but not individual stem cells. Blood 111(12):5553–5561.

21. Dykstra B, Olthof S, Schreuder J, Ritsema M, de Haan G (2011) Clonal analysis reveals multiple functional defects of aged murine hematopoietic stem cells. J Exp Med:jem. 20111490.

22. Muller-Sieburg CE, Sieburg HB, Bernitz JM, Cattarossi G (2012) Stem cell heterogeneity: implications for aging and regenerative medicine. Blood 119(17):3900–3907.

23. Rossi DJ, Jamieson CH, Weissman IL (2008) Stems cells and the pathways to aging and cancer. Cell 132(4):681–696.

24. Purton LE, Scadden DT (2007) Limiting factors in murine hematopoietic stem cell assays. Cell Stem Cell 1(3):263–270.

25. Domen J, Gandy KL, Weissman IL (1998) Systemic overexpression of BCL-2 in the hematopoietic system protects transgenic mice from the consequences of lethal irradiation. Blood 91(7):2272–2282.

26. Czechowicz A, Kraft D, Weissman IL, Bhattacharya D (2007) Efficient transplantation via antibody-based clearance of hematopoietic stem cell niches. Science 318(5854):1296–1299.

27. Dominici M, et al. (2009) Restoration and reversible expansion of the osteoblastic hematopoietic stem cell niche after marrow radioablation. Blood 114(11):2333–2343.

28. Pietras EM, et al. (2015) Functionally distinct subsets of lineage-biased multipotent progenitors control blood production in normal and regenerative conditions. Cell Stem Cell 17(1):35–46.

29. Allsopp RC, Cheshier S, Weissman IL (2001) Telomere shortening accompanies increased cell cycle activity during serial transplantation of hematopoietic stem cells. J Exp Med 193(8):917–924.

30. Ito M, et al. (2005) Stem cells in the hair follicle bulge contribute to wound repair but not to homeostasis of the epidermis. Nat Med 11(12):1351–1354.

31. Tian H, et al. (2011) A reserve stem cell population in small intestine renders Lgr5-positive cells dispensable. Nature 478(7368):255–259.

32. Bhattacharya D, et al. (2009) Niche recycling through division-independent egress of hematopoietic stem cells. J Exp Med 206(12):2837–2850.

33. Goodman JW, Hodgson GS (1962) Evidence for stem cells in the peripheral blood of mice. Blood 19(6):702–714.

34. McCredie KB, Hersh EM, Freireich EJ (1971) Cells capable of colony formation in the peripheral blood of man. Science 171(3968):293–294.

35. Wright DE, Wagers AJ, Gulati AP, Johnson FL, Weissman IL (2001) Physiological migration of hematopoietic stem and progenitor cells. Science 294(5548):1933–1936.

36. Lu R, Neff NF, Quake SR, Weissman IL (2011) Tracking single hematopoietic stem cells in vivo using high-throughput sequencing in conjunction with viral genetic barcoding. Nat Biotechnol 29(10):928–933.

37. Brewer C, Chu E, Chin M, Lu R (2016) Transplantation dose alters the differentiation program of hematopoietic stem cells. Cell Rep 15(8):1848–1857.

38. Nguyen L, et al. (2017) Functional Compensation Between Hematopoietic Stem Cells In Vivo. bioRxiv. doi:10.1101/236398.

39. Morrison SJ, Weissman IL (1994) The long-term repopulating subset of hematopoietic stem cells is deterministic and isolatable by phenotype. Immunity 1(8):661–673.

40. Yang L, et al. (2005) Identification of Lin-Sca1+ kit+ CD34+ Flt3-short-term hematopoietic stem cells capable of rapidly reconstituting and rescuing myeloablated transplant recipients. Blood 105(7):2717–2723.

41. Fialkow PJ, Jacobson RJ, Papayannopoulou T (1977) Chronic myelocytic leukemia: clonal origin in a stem cell common to the granulocyte, erythrocyte, platelet and monocyte/macrophage. Am J Med 63(1):125–130.

42. Serwold T, Ehrlich LIR, Weissman IL (2009) Reductive isolation from bone marrow and blood implicates common lymphoid progenitors as the major source of thymopoiesis. Blood 113(4):807–815.

43. Beachy PA, Karhadkar SS, Berman DM (2004) Tissue repair and stem cell renewal in carcinogenesis. Nature 432(7015):324–331.

44. Cartier N, et al. (2009) Hematopoietic stem cell gene therapy with a lentiviral vector in X-linked adrenoleukodystrophy. science 326(5954):818–823.

45. Cavazzana-Calvo M, et al. (2010) Transfusion independence and HMGA2 activation after gene therapy of human [bgr]-thalassaemia. Nature 467(7313):318–322.

46. Gerrits A, et al. (2010) Cellular barcoding tool for clonal analysis in the hematopoietic system. Blood 115(13):2610–2618.

47. Jordan CT, Lemischka IR (1990) Clonal and systemic analysis of long-term hematopoiesis in the mouse. Genes Dev 4(2):220–232.

48. Lemischka IR, Raulet DH, Mulligan RC (1986) Developmental potential and dynamic behavior of hematopoietic stem cells. Cell 45(6):917–927.

49. Wilson A, et al. (2008) Hematopoietic stem cells reversibly switch from dormancy to self-renewal during homeostasis and repair. Cell 135(6):1118–1129.

50. Prchal JT, et al. (1996) Clonal stability of blood cell lineages indicated by X-chromosomal transcriptional polymorphism. J Exp Med 183(2):561–567.

51. Orkin SH (2000) Diversification of haematopoietic stem cells to specific lineages. Nat Rev Genet 1(1):57–64.

52. Hsieh MM, et al. (2009) Allogeneic hematopoietic stem-cell transplantation for sickle cell disease. N Engl J Med 361(24):2309–2317.

53. Maloney DG, Sandmaier BM, Mackinnon S, Shizuru JA (2002) Non-myeloablative transplantation. ASH Educ Program Book 2002(1):392–421.

54. Challen GA, Boles NC, Chambers SM, Goodell MA (2010) Distinct hematopoietic stem cell subtypes are differentially regulated by TGF-β1. Cell Stem Cell 6(3):265–278.

55. Kent DG, et al. (2009) Prospective isolation and molecular characterization of hematopoietic stem cells with durable self-renewal potential. Blood 113(25):6342–6350.

